# Chemoselective Characterization of New Extracellular Matrix Deposition in Bioengineered Tumor Tissues

**DOI:** 10.1101/2025.03.18.643336

**Authors:** Zihan Ling, Burke Niego, Qingyang Li, Vanessa Serna Villa, Dhruv Bhattaram, Michael Hu, Zhuowei Gong, Lloyd M. Smith, Brian L. Frey, Xi Ren

## Abstract

The extracellular matrix (ECM), present in nearly all tissues, provides extensive support to resident cells through structural, biomechanical, and biochemical means, and in return the ECM undergoes constant remodeling from interacting cells to adapt to the evolving tissue states. Bioengineered 3D tissues, commonly known as cell-ECM composites, are robust model systems to recapitulate and investigate native pathophysiology. Key to this engineered morphogenesis process are the intricate cell-ECM interactions reflected by how cells respond to and thereby modulate their surrounding microenvironments through their ongoing ECM secretome. However, investigating ECM-regulated new ECM production has been challenging due to the proteomic background from the pre-existing biomaterial ECM. To address this hindrance, here we present a chemoselective strategy to label, enrich, and characterize newly synthesized ECM (newsECM) proteins produced by resident cells, allowing distinction from the pre-existing ECM background. Applying our analytical pipeline to bioengineered tumor tissues, either built upon decellularized ECM (dECM-tumors) or as ECM-free tumor spheroids (tumoroids), we observed distinct ECM synthesis patterns that were linked to their extracellular environments. Tumor cells responded to the dECM presence with elevated ECM remodeling activities, mediated by augmented digestion of pre-existing ECM coupled with upregulated synthesis of tumor-associated ECM. Our findings highlight the sensitivity of newsECM profiling to capture remodeling events that are otherwise under-represented by bulk proteomics and underscore the significance of dECM support for enabling native-like tumor cell behaviors. We anticipate the described newsECM analytical pipeline to be broadly applicable to other tissue-engineered systems to probe ECM-regulated ECM synthesis and remodeling, both fundamental aspects of cell-ECM crosstalk in engineered tissue morphogenesis.

## Introduction

Constructed with proteins and glycosaminoglycans as main building blocks, the extracellular matrix (ECM) is a complex meshwork that not only provides structural support to resident cells but also acts as a signaling pool, delivering critical microenvironmental cues to regulate cellular pathophysiology^[1,2]^. Contemporaneously, the interaction between the ECM and cells is not unidirectional. Instead of remaining static, the ECM undergoes dynamic remodeling by resident cells through processes such as deposition, modification, and degradation ^[3]^. Frequently, ECM remodeling is observed to progress in synergy with cellular transformation to mediate both physiological tissue morphogenesis as well as pathological fibrogenesis and tumorigenesis ^[3,4]^. A key mechanism for cells to remodel their surrounding ECM is through regulated secretion of new ECM components, in particular proteins, into the extracellular space that either make a structural contribution to the formation of new ECM structures or modify/degrade the pre-existing ECM ^[5]^. Thus, capturing and understanding the dynamic synthesis and deposition of new ECM proteins is essential to elucidate how resident cells respond to their evolving cellular and extracellular states in various pathophysiological contexts.

Tumorigenesis is characterized by somatic cell transformation into a state characterized by uncontrolled proliferation, with a further potential to invade and spread into other tissues ^[6–8]^. The unique ECM properties of solid tumors have long been recognized with generally elevated stiffness and vascularization compared to healthy tissue counterparts, favoring cancer cell survival and expansion. The acquisition of these tumor-specific ECM features is a result of continuous ECM remodeling by the tumor cells ^[9]^. The tumor cells secrete proteases such as Matrix Metalloproteinases (MMPs) which degrade the original healthy ECM, replacing it with tumor ECM ^[9]^. They also release signaling proteins such as growth factors to the extracellular niche to promote angiogenesis and metastasis ^[8]^.

Bioengineered 3D tissues have emerged as powerful model systems for recapitulating disease processes, investigating pathogenic mechanisms, and developing new therapeutics ^[10–12]^. A common strategy for engineering 3D tissue is to combine cells with ECM materials that deliver the desired biochemical composition, biomechanical property, and/or architecture of tissue-specific extracellular microenvironment; or a combination of all aspects, as occurs with decellularized native organ scaffolds ^[13]^. Nonetheless, it is also observed that in the absence of exogenous ECM support many cell types, such as benign epithelial cells and tumor cells, have the intrinsic capability to self-assemble into organotypic structures in the form of spheroids or organoids ^[14,15]^, where the extracellular architecture is formed solely by *de novo* ECM synthesis and deposition of the constituent cells. How do cells remodel the ECM materials that they are attached to, and how does the presence of pre-existing ECM modulate the ECM secretome of seeded cells? These fundamental questions in tissue engineering are yet to be fully understood.

A key challenge in studying the ECM-modulation of new ECM production in the context of tissue engineering is how to differentiate new versus pre-existing ECM proteins. Stable isotopic labeling of amino acids in cell culture (SILAC) ^[16]^ has enabled the proteomic detection of new protein synthesis through the incorporation of heavy-isotope amino acids ^[17]^. However, mass spectrometry-based proteomics is an abundance-proportional technique that favors identifying high-abundance protein species, but it falls short in capturing subtle changes of low-abundance regulatory proteins that can play key roles in altering the ECM microenvironment through direct (crosslinking, chemical modification, and degradation) and indirect (modulating cellular behavior and their ECM secretome) mechanisms ^[18]^. Other technologies, such as RNA sequencing, only allow us to access the instantaneous ECM gene expression at the transcriptomic level ^[19]^. Therefore, there is a critical need for a novel proteomics-based technology that reveals information related to translational and post-translational control, which ultimately determines the quantity, stability, and biochemical properties of newly deposited ECM proteins ^[20,21]^.

Glycosylation is one of the most common types of post-translational modifications (PTMs) where polysaccharide side chains are added to the newly synthesized core proteins. Glycosylation takes place primarily in the endoplasmic reticulum (ER) and Golgi apparatus, which are the organelles that synthesize, modify, and transport proteins that are destined for secretion into the extracellular space. Therefore, glycosylation is highly prevalent in ECM and ECM-associated proteins ^[22]^; in fact, it is estimated to occur for more than 90% of secreted proteins in humans ^[23–25]^. Consistent with this, our prior work has established the high efficacy of bioorthogonal labeling and detection of new ECM deposition using azido-tag-bearing monosaccharide probes both in vivo and ex vivo, with azido-galactosamine tetra-acetylated (Ac_4_GalNAz) being the most effective ^[26,27]^. These azido-tagged ECM proteins are compatible with chemoselective conjugation via click chemistry with a wide variety of ligands that enable a broad spectrum of downstream applications such as ECM material functionalization with bioactive molecules ^[28]^.

In this study, we navigated bioorthogonal ECM labeling to a new direction for ECM discovery and established a chemoselective strategy to profile newly synthesized ECM (newsECM) produced by resident cells and secreted into the pre-existing ECM scaffolds. Using a bioengineered lung cancer model combining tumor cells and acellular whole-lung scaffold, we demonstrated effective labeling, detection, enrichment, and proteomic identification of newly synthesized ECM (newsECM) glycoproteins. Leveraging this robust analytical pipeline, we characterized and compared the newsECM profiles within tumor tissues derived from the same tumor cells and formed with or without exogenous ECM scaffolds. Our comparison revealed elevated ECM remodeling activity of the tumor cells in the presence of pre-existing ECM. We anticipate the presented technology to be widely applicable for investigating cell-ECM interactions and ECM-regulated ECM remodeling in the context of tissue engineering.

## Methods

### Cell Culture

The human bronchioalveolar carcinoma cell line NCI-H358 (ATCC, CRL-5807) was grown as adherent cultures in complete medium composed of RPMI-1640 supplemented with 10% fetal bovine serum (FBS). All cells used in this study tested negative for mycoplasma.

### Whole Lung Perfusion Decellularization

All animal experiments were approved by and performed according to the guidelines of the Institutional Animal Care and Use Committee (IACUC) at Carnegie Mellon University. Heart-lung blocs were isolated from Sprague-Dawley rats (200-250g, Charles River Laboratories, Strain Code 400), cannulated and perfusion decellularized as previously described ^[29]^. Briefly, rats were euthanized under CO_2_ inhalation, followed by a flushing of PBS through the pulmonary artery (PA) and the isolation of heart-lung bloc. The PA is then cannulated and perfusion decellularized with 0.1% SDS, followed by sequential washing of water, 1% Triton X-100 and PBS. The decellularized (dECM) lung scaffolds were preserved in sterile DPBS supplemented with anti-biotic/anti-mycotic at 4°C.

### Establishment of the dECM-tumor model and Metabolic Labeling of Newly Synthesized Proteins

8 dECM lung scaffolds were randomly assigned into 2 groups: 5 for the Ac_4_GalNAz group and 3 for vehicle (DMSO) control group, differentiated by the chemical administration on day 6. The dECM lung scaffolds were perfusion washed with PBS for 30 minutes before the seeding procedure. 50 million NCI-H358 cells were collected from culture flasks and resuspended to 50 mL. The dECM scaffolds were flushed with 30 mL DPBS through the trachea under approximately 40 cmH_2_O gravity pressure and then seeded with 50 million resuspended cells through the trachea under the same gravity pressure ^[30]^. The trachea was ligated after the instillation to maintain airway pressure. The dECM-tumor was perfusion cultured with complete medium (RPMI-1640 supplemented with 10% FBS, 100 units/mL of penicillin and 100 μg/mL of streptomycin) with the perfusion rate at 1.26 mL/min in the first 24 hours, and 5.04 mL/min in the following 6 days. A complete medium change was performed every 2 days. On day 6, the medium was changed to complete medium supplemented with Ac_4_GalNAz (Vector Laboratories, CCT-1086; final concentration 50 µM with 0.1% DMSO) or DMSO (final concentration 0.1%, as vehicle control) and the dECM-tumors were harvested 24 hours later on day 7. Upon harvesting, the dECM-tumors were flushed with 10 mL DPBS through PA to remove the leftover medium in the scaffold. Then the left lobe and right lower lobe were ligated and removed for protein extraction. The remaining lobes were gravity-fixed with 4% PFA in PBS for histological analysis.

### Generating NCI-H358 Tumoroids and Metabolic Labeling of Newly Synthesized Proteins

3.2 million NCI-H358 cells were resuspended in 4 mL complete medium (RPMI-1640 supplemented with 10% FBS, 100 units/mL of penicillin and 100 μg/mL of streptomycin) into each well of a 6-well cell-repellent plate. Every 2 wells of tumoroids were considered as 1 sample for subsequent analysis and 10 samples (from 20 wells) were randomly assigned into 2 groups: 5 samples (from 10 wells) for the Ac_4_GalNAz group and 5 samples (from 10 wells) for vehicle (DMSO) control group, differentiated by the chemical administration on day 6. The plates were placed on an orbital shaker at 125 rpm with half of the medium changed daily for tumoroid formation and maintenance. On day 6, the medium was changed to complete medium supplemented with Ac_4_GalNAz (final concentration 50 µM with 0.1% DMSO) or DMSO (final concentration 0.1%, as vehicle control). After 24 hours, the tumoroids were collected and washed with DPBS, followed by protein extraction or fixation under 4% PFA in PBS. The fixed tumoroids were embedded in HistoGel for histological analysis.

### Histology and Immunostaining

The fixed dECM-tumors or HistoGel-embedded tumoroids were sectioned and stained with hematoxylin and eosin (H&E). For immunofluorescence staining, the sections were incubated for antigen unmasking in citrate-based solution and permeabilized with 0.1% Triton X-100. For immunofluorescence detection of azido signal, the sections were reacted to Alkyne-PEG4-Biotin (Vector Laboratories, CCT-TA105) under copper-catalyzed cycloaddition (CuAAC) condition ^[31]^. The sections were blocked with 1% bovine serum albumin (BSA) in DPBS and stained with streptavidin conjugated to Alexa Fluor-647 (Thermo Fisher Scientific, S21374) at the dilution of 1:500. For antibody-based immunofluorescence staining, the sections were blocked with 1% BSA in DPBS and incubated with primary antibody for MMP-14 (Abcam, ab51074) at the dilution of 1:100, E-Cadherin (Cell Signaling Technology, 14472) at the dilution of 1:200, human Laminin Subunit Gamma 2 (hLAMC2, Abcam, ab210959) at the dilution of 1:500 or Laminin Subunit Alpha 1 (LAMA1, Abcam, ab11575) at the dilution of 1:500. The sections were stained with secondary antibodies conjugated to Alexa Fluor-647 or Alexa Fluor-488 at the dilution of 1:500. All slides were mounted with DAPI-containing Fluoromount solution (SouthernBiotech, 0100-20).

### Immunofluorescence Quantification

Immunofluorescence images were taken under EVOS FL Auto 2 Imaging System (Thermo Fisher Scientific) or Nikon A1R HD25 Confocal Microscope System (Nikon Instruments Inc.). For dECM-tumor sections, five random fields were taken for each biological replicate; for tumoroid sections, each tumoroid was considered as a replicate. Fluorescent images were split into their component RGB channels. The blue channel, containing the DAPI signal in all images taken, was converted into a map of the tissue within the image via a custom MATLAB program. The program adjusts the saturation to balance the blue channel before being passed through a threshold equal to 20% of maximum pixel intensity to filter out noise, binarizing the image in the process. Size filtration was then applied to the map to remove non-nuclei debris. The identified nuclei blobs were then dilated and eroded by a disk structuring element to connect the blobs in a manner that approximates the underlying tissue, resulting in a “DAPI map”. Finally, the average pixel intensities (normalized to scale of 0 to 1) of the red and green channels (each representing stains for various proteins) both within and outside the approximated tissue area were calculated through logical indexing via this DAPI map. The resulting intensities from red or green channels were then used to calculate normalized immunofluorescence intensities. All immunofluorescence images presented in figures were processed by Fiji software ^[32]^.

### Stepwise Extraction of Cellular and ECM Protein Fractions and Biotinylation

For the dECM-tumors, the tissues were minced into fine pieces with surgical scissors and homogenized in 2-mL tubes prefilled with glass beads (Benchmark Scientific, D1031-10). The homogenized tissues were washed with DPBS to remove the perfusate residues. The cellular fractions were extracted at 4°C with 8 mM 3-[(3-cholamidopropyl) dimethylammonio]-1-propanesulfonate (CHAPS, Thermo Fisher Scientific, 28300) buffer prepared in DPBS (pH 5.50) that is supplemented with 1 M NaCl and 1% protease and phosphatase inhibitor cocktail (Thermo Fisher Scientific, 78440). The insoluble pellets were washed with DPBS, and the ECM fractions were extracted with 4M Urea (Millipore Sigma, U5128), 0.1% SDS in DPBS supplemented with 1% protease and phosphatase inhibitor cocktail. For the ECM-free tumoroids, the samples were incubated at 4°C under agitation with 8 mM CHAPS buffer prepared in DPBS (pH 5.50) supplemented with 1 M NaCl and 1% protease and phosphatase inhibitor cocktail to extract cellular fractions. The insoluble pellets were then washed with DPBS, and the ECM fractions were extracted with 4 M Urea, 0.1% SDS in DPBS supplemented with 1% protease and phosphatase inhibitor cocktail. The extracts were reacted to Alkyne-PEG4-biotin under the CuAAC condition for Western blot analysis.

### Western Blots and In-Gel/Dot blot Total Protein Staining

Western blotting and in-gel total protein staining were performed as previously described ^[27]^. Briefly, the protein samples were quantified with bicinchoninic acid (BCA) assay, and an equal amount of protein was analyzed using sodium dodecyl-sulfate polyacrylamide gel electrophoresis (SDS-PAGE). For Western blot, the proteins in gel were transferred to polyvinylidene difluoride (PVDF) membranes, incubated with primary antibodies for MMP-14 (Abcam, ab51074) at the dilution of 1:1000, GAPDH (Santa Cruz Biotechnology, sc-32233) at the dilution of 1:500 or Laminin Subunit Alpha 1 (LAMA1, Abcam, ab11575) at the dilution of 1:2000, followed by staining with secondary antibodies conjugated to horseradish peroxidase (HRP) for autoradiography. For in-gel total protein staining, PAGE gels were fixed with 50% methanol and 7% acetic acid, stained with SYPRO Ruby protein gel stain (Thermo Fisher Scientific, S12000) and visualized under ultraviolet light. For dot blot total protein stain, 3 μL of protein samples were applied to nitrocellulose membranes. The membranes were incubated in 10% methanol, 7% acetic acid, stained with SYPRO Ruby protein blot stain (Thermo Fisher Scientific, S11791) and visualized under ultraviolet light. Image colors of SYPRO ruby stains were inverted. Signal intensities were measured with Fiji software.

### Chemoselective Enrichment of Azido-Tagged Proteins

The ECM extracts from dECM-tumors and tumoroids were reacted with Alkyne-PEG4-Desthiobiotin (Vector Laboratories, CCT-1109) under the CuAAC condition. The unreacted Alkyne-PEG4-Desthiobiotin was removed with buffer exchange into 5 M Urea, 0.1% SDS in DPBS (pH 7.40) using a 3 kDa molecular weight cut-off filter (Millipore Sigma, UFC5003), and the retentates were saved as “input” samples. Following quantification by Pierce^TM^ BCA protein assay kit (Thermo Fisher Scientific, 23225), aliquots of each input sample containing 1 mg of protein (with a final volume of 1.5 mL) were introduced to 50 µL suspending streptavidin-conjugated resins (Thermo Fisher Scientific, 53116) for pull-down of desthiobiotinylated proteins. Following 5-hour binding, the supernatants were collected, and the resins were washed 8 times, 15 min per time, with 5 M Urea, 0.1% SDS in DPBS (pH 7.40). The pulled-down proteins were eluted with 500 μL of 5 M Urea, 0.1% SDS in DPBS (pH 7.40) supplemented with 30 mM biotin at room temperature for 1 h. The resulting eluates were concentrated with a 10 kDa molecular weight cut-off filter (Millipore Sigma, UFC5010) to 90 μL. 18 μL of each “eluate” sample was used for SDS-PAGE to test the enrichment efficacy and specificity; the remaining 72 μL was stored at -80°C.

### Liquid Chromatography-Tandem Mass Spectrometry Analysis

The 72 μL “eluate” samples were thawed for proteomic characterization. Analogous “input” samples were prepared by diluting aliquots of each input sample containing ∼3.6 μg of protein (based upon BCA results) to 72 μL with 5 M Urea, 0.1% SDS, in PBS pH 7.40. Disulfide bonds were reduced with 5 mM dithiothreitol (DTT, Millipore Sigma) and heated to 37 °C for one hour; after cooling, samples were alkylated in the dark for 30 min using 30 mM iodoacetamide (Millipore Sigma). Alkylation was quenched with 5 mM DTT at room temperature for 15 min. Samples were prepared for mass spectrometry using Single-Pot, Solid-Phase-enhanced Sample Preparation (SP3) ^[33]^. Briefly, samples were reconstituted by adding 100 μg of Sera-Mag^TM^ carboxylate-modified magnetic beads (50 μg/μL, Cytiva) to each sample, followed by an equal volume (90.6 μL) of absolute ethanol (Millipore Sigma). The samples were mixed at room temperature for 5 min on a thermomixer at 1,000 rpm to allow protein binding. Afterwards, samples were placed on a magnetic stand for 2 min and the supernatant was removed. Beads were washed three times by sequential resuspension in 180 μL of 80% ethanol, placing on a magnetic stand for 2 min, and removing the supernatant. Proteolytic digestion occurred on the beads after resuspension in 100 μL of 50 mM ammonium bicarbonate (ABC, Millipore Sigma) with 0.14 μg of trypsin (Promega, 1:25 ratio) for the inputs and dECM-tumor eluates and 0.072 μg of trypsin for the tumoroid eluates. The samples were digested for 2 hours at 47 °C, 1000 rpm. Following digestion, the samples were centrifuged at 20,000xg for 1 min at room temperature. The supernatant containing the tryptic peptides was removed, transferred to a new tube, and evaporated to dryness. Samples were reconstituted in 150 μL of 0.1% trifluoracetic acid (TFA) and cleaned up using a C18 ZipTip (Pierce, Thermo Fisher Scientific, 87784) according to the manufacturer’s instructions. After evaporation to dryness, samples were reconstituted with 8 μL of 5% acetonitrile containing 0.2% formic acid. Liquid chromatography-tandem mass spectrometry analysis was performed with 4 μL of each eluate sample and 2 μL of each input sample.

Samples were analyzed by a HPLC-MS/MS system consisting of a high-performance liquid chromatograph (HPLC, nanoAcquity, Waters) connected to an electrospray ionization (ESI) Orbitrap mass spectrometer (QE HF, Thermo Fisher Scientific). Injected peptides were loaded onto a 20 cm long fused silica capillary nano-column packed with C18 beads (1.7 μm in diameter, 130 Angstrom pore size from Waters BEH), with an emitter tip pulled approximately to 1 μm using a laser puller (Sutter instruments). Peptides were initially loaded on-column for 30 minutes at 400 nL/min at 98% buffer A (aqueous 0.1% formic acid) and 2% buffer B (acetonitrile with 0.1% formic acid) and then eluted over 120 min at a flow rate of 300 nL/min with a gradient as follows: time 1 min-2% buffer B; time 31 min-8% buffer B; time 111 min-44% buffer B; time 122-129 min-64% buffer B; time 133-152 min-equilibrate at 2% buffer B. The nano-column was held at 60 °C using a column heater constructed in house.

The nanospray source voltage was set to 2,200 V. Full-mass profile scans were performed in the FT-orbitrap between 375-1500 m/z at a resolution of 120,000, followed by MS/MS higher-energy collisional dissociation (HCD) scans of the ten highest intensity parent ions at 30% collision energy (CE) and 15,000 resolution, with a mass range starting at 100 m/z and a 2.5 m/z isolation window. Charge states 2-5 were included and dynamic exclusion was enabled with a repeat count of one over a duration of 15 s. The MS/MS orbitrap HCD scans were collected with automatic gain control (AGC) target set to 1e5 ions, a maximum inject time of 500 ms, and 1 microscan per spectrum.

### Proteomic Data Analysis

LC-MS/MS data were analyzed with the open search software program MetaMorpheus (version 1.0.5, available at https://github.com/smith-chem-wisc/MetaMorpheus). The Swiss-Prot Rat XML (reviewed) database containing 8,207 protein entries [downloaded from UniProt (UniProtKB, 10116) 04/01/2024] and the Swiss-Prot Human XML (reviewed) database containing 20,435 protein entries [downloaded from UniProt (UniProtKB, 9606) 04/01/2024] were utilized along with the MetaMorpheus default contaminants database. MetaMorpheus was used to calibrate the raw data files for the 36 samples (18 inputs and 18 eluates) and then subject the files to global posttranslational modification (PTM) discovery (GPTMD) ^[34]^ to identify possible PTMs not already annotated in the employed databases. The modifications searched included: the Common Biological, Common Artifact, and Metal Adduct categories within MetaMorpheus, as well as custom modifications for the azido-glycan labels (on serine/threonine with and without the alkyne-PEG4-desthiobiotin) at +244.0808 Da and +671.3490 Da. The files were then searched against the GPTMD databases using the following search parameters: protease = trypsin; search for truncated proteins and proteolysis products = False; maximum missed cleavages = 2; minimum peptide length = 7; maximum peptide length = unspecified; initiator methionine behavior = Variable; fixed modifications = Carbamidomethyl on C, Carbamidomethyl on U; variable modifications = Oxidation on M; max mods per peptide = 2; max modification isoforms = 1024; precursor mass tolerance = ±5.0000 ppm; product mass tolerance = ±20.0000 ppm; report PSM ambiguity = True.

Using the above parameters, the following three searches were completed in MetaMorpheus with label-free quantification (LFQ) enabled: (1) all 18 eluate files underwent GPTMD and were searched using the Human XML database with Match-Between-Runs (MBR), (2) all 16 dECM-tumor files (inputs and eluates) underwent GPTMD analysis and were searched using the Human and Rat XML databases without MBR, and (3) the 18 dECM-tumor and tumoroid input files underwent GPTMD analysis and were searched using the Human and Rat XML databases with MBR.

Data analysis details for *Search 1* (all 18 eluates) are as follows. All 18 eluates files underwent GPTMD and were searched using the Human XML database with MBR. A one-way ANOVA was done on protein IDs from each sample across 4 treatment groups (Ac_4_GalNAz, dECM-tumor; Ac_4_GalNAz, tumoroid; vehicle, dECM-tumor; vehicle, tumoroid) and outliers were removed for subsequent analyses. The intensities of all human proteins were summed for each sample. For comparison between Ac_4_GalNAz and vehicle groups, the protein intensities were normalized separately in each treatment group. The normalized intensity (Table 2 – “*Search 1* normalization 1”) of protein *X* in sample *i* under treatment group *K* (*K* can be one of the following: Ac_4_GalNAz, dECM-tumor; Ac_4_GalNAz, tumoroid; vehicle, dECM-tumor; vehicle, tumoroid) was calculated with the following formula:

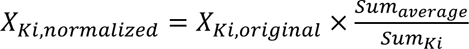, where *Sum_average_* was the average sum of all eluate samples in treatment group *K*.

The dataset was then separated into two subsets based on their culture models (dECM-tumor or tumoroid) for subsequent analysis. Using Perseus ^[35]^ each subset was pre-filtered to remove proteins with less than two nonzero values in each treatment group. Analysis in Perseus continued with the filtered list of protein intensities by first performing a Log_2_ transform and then undergoing a two-tailed Welch’s *t*-test using criteria in the Statistical Analysis section below.

Another comparison from the *Search 1* dataset was between Ac_4_GalNAz dECM-tumor and Ac_4_GalNAz tumoroid groups. Proteins from these 9 samples were normalized all together for their proportional intensities in each group. The normalized intensity (Table 2 – “*Search 1* normalization 2”) of protein *X* in sample *j* was calculated with the following formula:

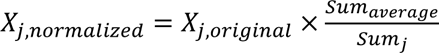, where *Sum_average_* was the average sum of all (both dECM-tumor and tumoroid) Ac_4_GalNAz eluate samples. The normalized intensities were used for Gene Ontology analysis. For volcano plots and Venn diagrams, the normalized protein intensities were taken as Log_2_ and imputed using the LMCD-QRILC R script plugin in Perseus ^[36,37]^. Briefly, the impute QRILC script imputed missing values by randomly drawing from a truncated normal distribution, that was estimated using quantile regression. Following imputation, the data underwent two-tailed Welch’s *t*-test using criteria in the Statistical Analysis section below. For individual protein intensity bar graphs, the imputed intensity values were transformed back to decimal numbers for plotting.

Data analysis details for *Search 2*: One eluate sample was removed as outlier, and the other 15 dECM-tumor files (inputs and eluates) underwent GPTMD analysis and were searched using the Human and Rat XML databases without MBR: Data was analyzed as original value (Table 2 – “*Search 2* original”) for calculations of total *Homo sapiens* and *Rattus Norvegicus* protein intensities.

Data analysis details for *Search 3*: All 18 tumoroid and dECM-tumor input files underwent GPTMD analysis and were searched using the Human and Rat XML databases with MBR: The intensities of all human and rat proteins were summed for each sample. All input samples were normalized together. The normalized intensity (Table 2 – “*Search 3* normalization”) of protein *X* in sample *m* was calculated with the following formula:

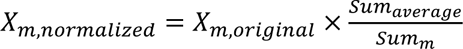, where *Sum_average_* was the average sum of all input samples. The normalized intensities were used for Gene Ontology analysis.

### Protein Composition and Kyoto Encyclopedia of Genes and Genomes (KEGG) Pathway Analysis

The KEGG pathway and protein composition analysis was performed and visualized on the Proteomaps website (https://www.proteomaps.net/index.html) ^[38]^. The protein list along with their normalized values were uploaded and analyzed for “*H. Sapiens*”.

### Gene Ontology Analysis

Gene ontology (GO) analysis was performed on the Database for Annotation, Visualization, and Integrated Discovery (DAVID) Bioinformatics Resources (https://david.ncifcrf.gov/tools.jsp) ^[39,40]^. The Uniprot ID lists were uploaded and analyzed for functional annotation clustering or enriched for biological process terms. The top annotation clusters with highest enrichment scores were presented in a table. The top GOTERM_BP_DIRECT terms with lowest *p* values were plotted in bar graphs.

### Protein-Protein Interaction Analysis

Protein-protein interaction analysis was performed and visualized with STRING database (https://string-db.org/) ^[41]^. Protein name was set as “MMP14”, Organisms was set as “*Homo sapiens*”, and Network type was set as “physical subnetwork”.

### Statistical Analysis

For all experiments, the *n* values stated represent the number of independent biological samples. Data was analyzed using GraphPad Prism, Microsoft Excel, RStudio, MATLAB, or Perseus. Statistical comparisons of data with three or more treatment groups were performed with one-way ANOVA with Tukey’s multiple comparisons tests (confidence level: 0.95). Between-group correlation coefficients were analyzed by Pearson correlation test. Outliers were identified and cleared out by Grubbs’s tests (*α* = 0.05). The multiple comparison testing of protein intensities for volcano plots and Venn diagrams were analyzed by two-tailed *t*-tests with Welch’s correction, followed by multiple hypothesis testing correction using a permutation-based false discovery rate (FDR) calculation. *q*-values were reported after permutation-based FDR correction; proteins with *q* < 0.05 (i.e. 5% FDR) were considered significant. For volcano plots, the Perseus output of - Log_10_(Welch’s *t*-test *p*-value) was plotted versus the fold changes (FCs) calculated from geometric means of the decimal protein intensities. Paired measurements were analyzed by paired *t*-tests. All other comparisons with two treatment groups were analyzed by two-tailed *t*-tests with Welch’s correction. Error bars were plotted by standard deviations (SD). All statistical significances were reported accordingly. ns, *p* > 0.05; \**p* < 0.05; \*\**p* < 0.01; \*\*\**p* < 0.001; ^#^*q* < 0.05.

### Schematics

All schematics were created with BioRender.com.

## Results

### Engineering 3D tumor models in the presence and absence of ECM support

Tumor cells are well known to extensively modulate their surrounding extracellular microenvironment to favor their expansion and metastasis ^[9]^. To model and investigate these key pathological processes, 3D tumor tissues can be engineered in vitro either with or without exogenous biomaterial support ^[30,42]^. Yet, understanding of how the pre-existing ECM environment modulates the ability of tumor cells to produce new ECM and the properties of these ECM depositions remain elusive. To investigate this, we established two 3D tumor tissue models using NCI-H358 lung cancer cells, either as cell-ECM composites or as ECM-free tumor spheroids (tumoroids). To engineer the cell-ECM composites, we leveraged decellularized ECM (dECM) scaffolds prepared from perfusion decellularization of whole rat lungs and seeded the cancer cells through intratracheal instillation into what was the epithelial compartment prior to decellularization (Figure 1A) ^[30]^. With perfusion culture through the well-preserved native pulmonary vascular bed ^[30,43]^, the seeded cancer cells effectively engrafted onto the dECM material, leading to formation of tumor node clusters by day 7 of culture (Figure 1B). These resulting engineered tumor tissues on dECM scaffolds were referred to as dECM-tumors. In parallel, we generated ECM-free tumoroids over the same culture duration through the self-assembly of cancer cells on a cell-repellent surface coupled with constant agitation (Figure 1F-H).

**Figure 1.**
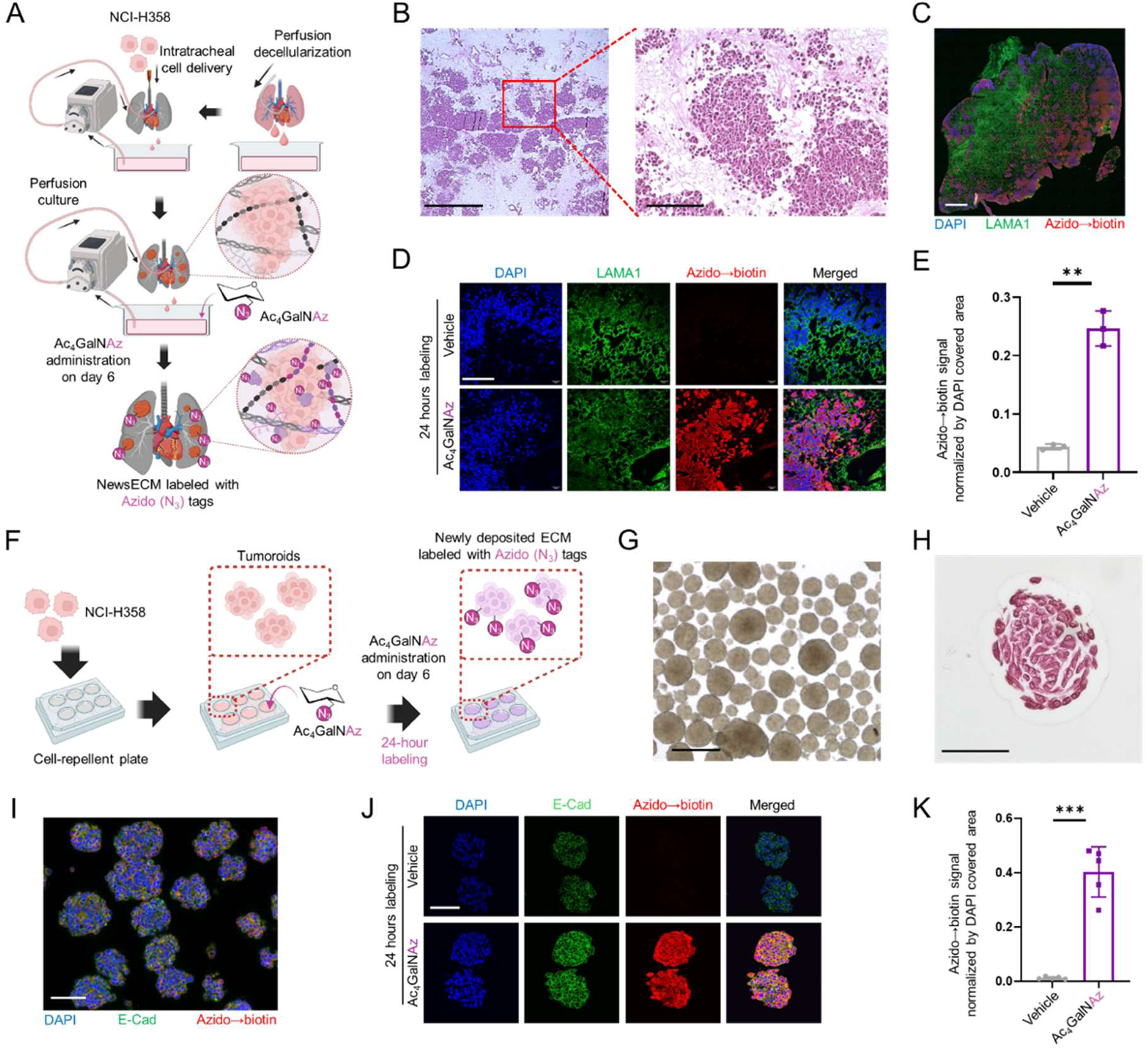
Glycosylation-enabled metabolic labeling of newsECM in engineered tumor models. (A) A schematic for the engineering of dECM-tumor model and metabolic newsECM labeling using Ac_4_GalNAz. (B) Hematoxylin and eosin (H&E) staining of the dECM-tumor after 7 days of perfusion culture. Scale bars: left, 1000 µm; right, 200 µm. (C,D) Stitched image (C) or single-field images (D) of immunofluorescence staining of azido→biotin (red) and LAMA1 (green) on dECM-tumors receiving Ac_4_GalNAz (*n*=3) or DMSO (vehicle control, *n*=3) during the last day of culture. Azido tags were visualized with fluorophore-conjugated streptavidin detection of biotin following alkyne-PEG4-biotin conjugation. Scale bar in panel C, 1000 µm. Scale bar in panel D, 100 µm. (E) Quantification of azido→biotin (red) signal intensity in panel D, normalized by DAPI covered area. (F) A schematic for generating the ECM-free tumoroid model and metabolic newsECM labeling using Ac_4_GalNAz. (G) Bright-field image of tumoroids at the end of 7-day culture. Scale bar, 500 µm. (H) H&E staining of tumoroids following 7 days of agitation culture. Scale bar: 75 µm. (I, J) Stitched image (I) or single-field images (J) of immunofluorescence staining of azido→biotin (red) and E-Cadherin (green) on tumoroid clusters receiving Ac_4_GalNAz (*n*=5) or DMSO (vehicle control, *n*=5) during the last day of culture. Scale bars in both panel I and panel J, 100 µm. (K) Quantification of azido→biotin (red) signal intensities in panel J, normalized by DAPI covered area. *** *p*<0.001; ** *p*<0.01. Data are presented as means ± SD. Schematics were created with Biorender.com and published with permission.

### Selective labeling of newsECM deposition in the dECM-tumor and ECM-free tumoroid using glycans with bioorthogonal tags

To selectively study the new ECM protein deposition by resident cells within a pre-existing ECM biomaterial environment, such as the dECM, we implemented a bioorthogonal labeling strategy that allows distinguishing newly synthesized ECM (referred to as newsECM) proteins from the pre-existing ones. Our strategy utilized an azido-tagged monosaccharide probe, Ac_4_GalNAz, that can be specifically incorporated into the glycans of newly synthesized glycoproteins, which include the majority of ECM proteins, during post-translational glycosylation ^[23–25]^. To apply this labeling procedure to the dECM-tumor model, we administered Ac_4_GalNAz (or DMSO as vehicle control) into the medium on day-6 of the culture and labeled for 24 hours (Figure 1A). The azido-tag incorporation was visualized through click conjugation with alkyne-biotin followed by fluorescence detection of biotin. We observed robust and specific azido incorporation within the dECM-tumor tissues, which further showed specific colocalization with regions of tumor nodule presence, which is consistent with the requirement of live cell metabolism for azido-probe incorporation into newly synthesized glycoproteins (Figure 1C-E). To allow subsequent comparison of newsECM profiles between dECM-tumors and ECM-free tumoroids, we performed a similar Ac_4_GalNAz labeling procedure on the tumoroids with the same duration (day-6 to day-7) and within the same medium; again, we observed specific azido incorporation throughout the tumoroids (Figure 1F,I-K).

### ECM extraction and chemoselective enrichment of newsECM

While the ECM is particularly enriched for glycoproteins, many cellular proteins are also glycosylated. To focus our analysis on the ECM, we developed a stepwise extraction method to separate the cellular from the extracellular protein contents. We first extracted the cellular protein fraction with a mild CHAPS buffer, followed by extraction of the extracellular protein fraction with a harsher urea buffer (Figure 2A). The extraction efficiency and specificity was validated by western blot analysis of cellular GAPDH or extracellular Laminin (Laminin Subunit Alpha 1, LAMA1), where we observed nearly exclusive presence of GAPDH and Laminin protein signals to the cellular and extracellular fractions, respectively, in both dECM-tumors and ECM-free tumoroids (Figure 2B,C left), with equal protein loading confirmed by SYPRO Ruby total protein stain (Figure 2B,C right). Further, we went on to examine azido labeling in each protein fraction via alkyne-biotin conjugation and on-blot biotin detection. Consistent with prior tissue staining findings, specific azido signals were detected in both cellular and ECM fractions of both dECM-tumors and tumoroids administered with Ac_4_GalNAz compared to those administered with vehicle control (Figure S1, Figure S2, Figure 2D,E).

**Figure 2.**
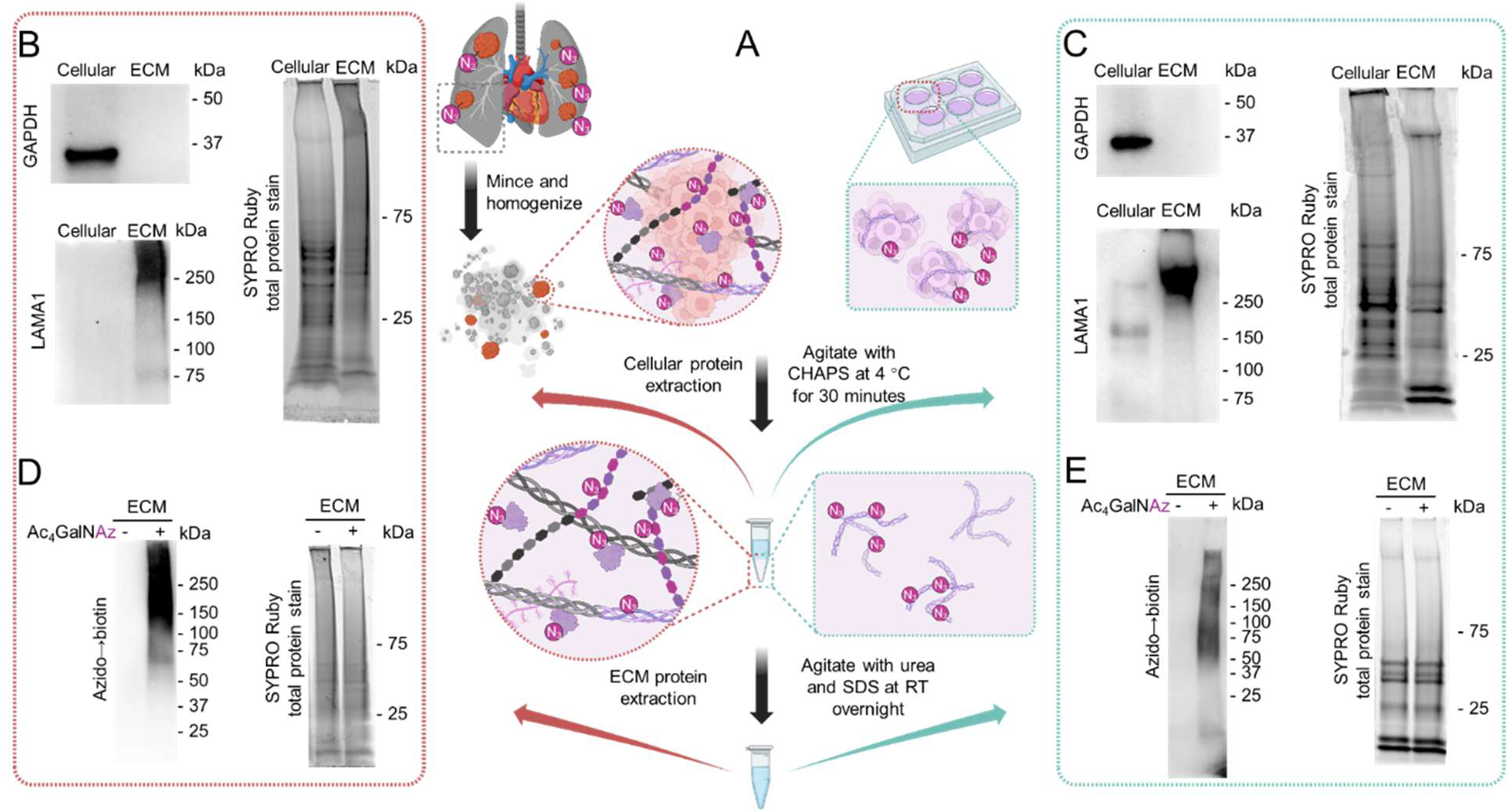
Stepwise extraction of cellular and ECM fractions in bioengineered tumor models. (A) Schematic for stepwise extraction of cellular and ECM proteins from dECM-tumors (left) and tumoroids (right). (B,C) Western blot detection of GAPDH (left, top) and LAMA1 (left, bottom) and SYPRO Ruby staining of total proteins (right) in cellular and ECM fractions from dECM-tumors (B) and tumoroids (C). (D,E) Western blot detection of azido→biotin signal in the ECM fractions of dECM-tumors (D) and tumoroids (E) using streptavidin-HRP (left) and SYPRO Ruby staining of total proteins (right).

To chemoselective purify the azido-tagged newsECM (Az-newsECM) from the total extracted ECM, we first converted azido labeling into desthiobiotin tags through click conjugation with alkyne-desthiobiotin and then performed streptavidin pull-down ^[27]^ (Figure 3A). We first demonstrated specific alkyne-desthiobiotin conjugation onto Az-newsECM from both dECM-tumors or ECM-free tumoroids with prior Ac_4_GalNAz administration (Figure 3B,C). Upon introducing the desthiobiotinylated ECM samples to streptavidin resins, we analyzed the azido→desthiobiotin signals from samples both before (“input”) and after (“supernatant”) incubation with streptavidin resins and observed a dramatic reduction in azido→desthiobiotin signals in the supernatants compared to inputs (Figure 3D,E), demonstrating high efficacy of streptavidin resins to capture desthiobiotinylated newsECM. Finally, we released the captured newsECM proteins through a competitive manner using free biotin with higher affinity to streptavidin compared to desthiobiotin ^[44]^ (Figure 3A). Further, SYPRO Ruby total protein stain confirmed specific protein elution from ECM inputs of Ac_4_GalNAz-labeled dECM-tumors and tumoroids, compared to elution from ECM inputs of tissues administered with vehicle control (Figure 3F,G). Altogether, these results demonstrate the effective and specific enrichment of Az-newsECM from engineered tumor tissues for subsequent proteomics analysis.

**Figure 3.**
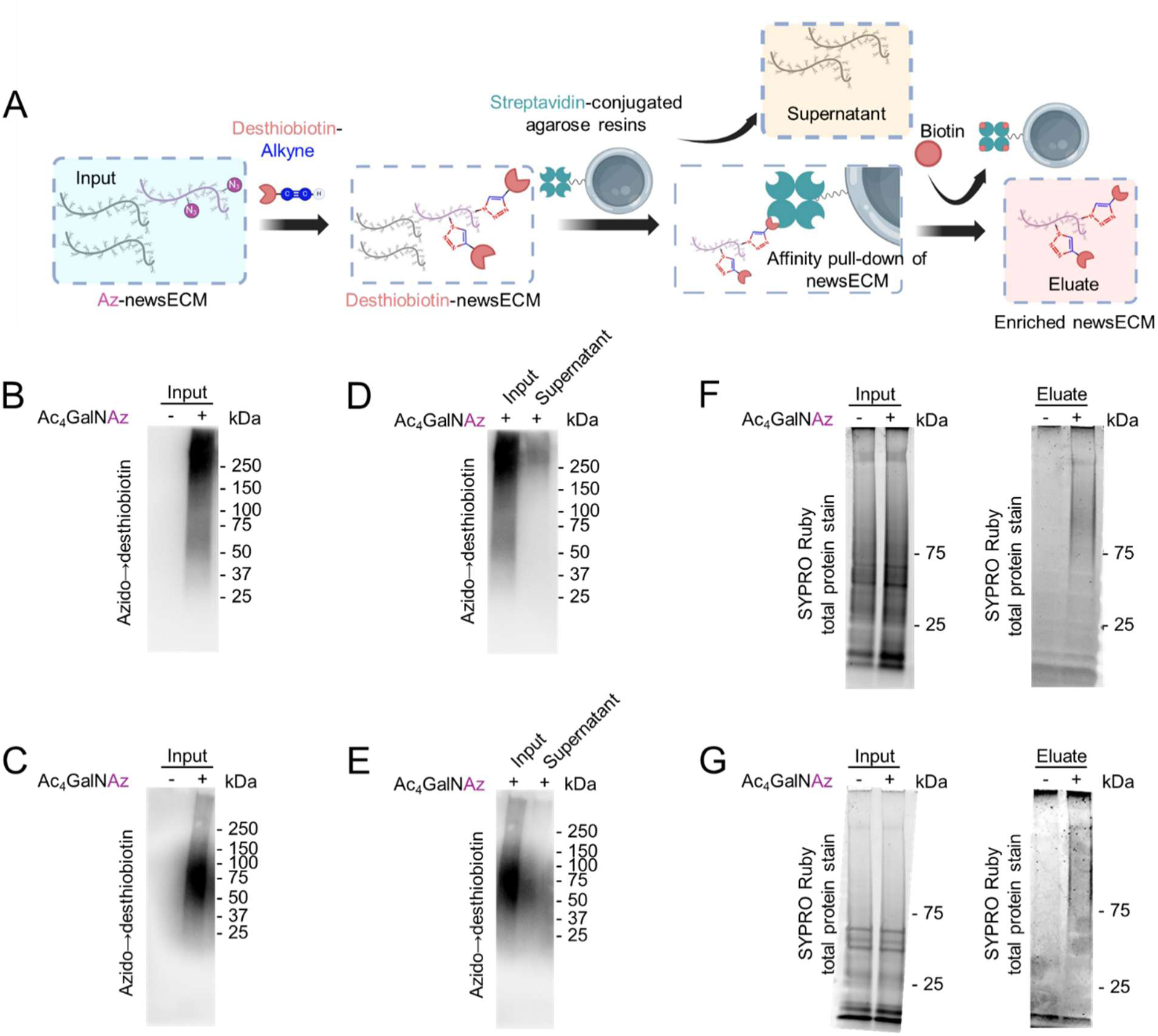
Chemoselective enrichment of azido-newsECM. (A) A schematic showing the workflow for chemoselective enrichment of Az-newsECM. The Az-newsECM were desthiobiotinylated (input), pulled down by streptavidin resins (the wash-offs were saved as supernatants), and released through competitive elution with high-affinity biotin (eluates). (B,C) Western blot analysis desthiobiotin signal in the ECM inputs of dECM-tumors (B) and tumoroids (C) receiving Ac_4_GalNAz or DMSO (vehicle control) using streptavidin-HRP. (D,E) Western blot analysis of desthiobiotin signal in the ECM inputs and supernatants of dECM-tumors (D) and tumoroids (E) receiving Ac_4_GalNAz using streptavidin-HRP. (F,G) SYPRO Ruby staining of total proteins in the ECM inputs (left) and eluates (right) of dECM-tumors (F) and tumoroids (G).

### Proteomics characterization of newsECM signatures

To characterize the newsECM signatures in both tumor models, we performed LC-MS/MS on the ECM samples both before (“inputs”) and after (“eluates”) the streptavidin pull-down (Figure 4A). Through the analysis, we observed significant augmentation in both the identified protein groups (5.3-fold augmentation in dECM-tumors and 9.0-fold augmentation in tumoroids) and total protein intensities (23.4-fold augmentation in dECM-tumors and 11.0-fold augmentation in tumoroids) in eluates from tumor tissues with prior administration of Ac_4_GalNAz probe compared to eluates from tissues exposed to vehicle control (Figure 4B,C). This observation is consistent with protein abundance results shown in the volcano plot comparisons (Figure 4D,E), the Venn diagrams (Figure 4F,G), the bar charts (Figure 4H,I) and with the total protein stain findings shown previously (Figure 3F,G); these results further validate our chemoselective enrichment approach for newsECM. Since the dECM-tumors were constructed combining human cells with dECM derived from rat lungs, we anticipate the newsECM profiling to be particularly enriched for human proteins that were directly labeled with azido tags but to a less extent to capturing rat proteins that can only be captured through indirect mechanisms such as interaction with the resin-bound Az-newsECM. To assess this, we normalized human- or rat-specific proteins identified in eluates based on their relative intensities in the inputs and observed that while human protein intensities detected in eluates remained at a level close to those of the inputs (with only 12.5% reduction), rat proteins in eluates were reduced by 70% compared to inputs (Figure S3). This finding supports the selectivity of the described newsECM profiling towards new protein production by metabolically active cells.

**Figure 4.**
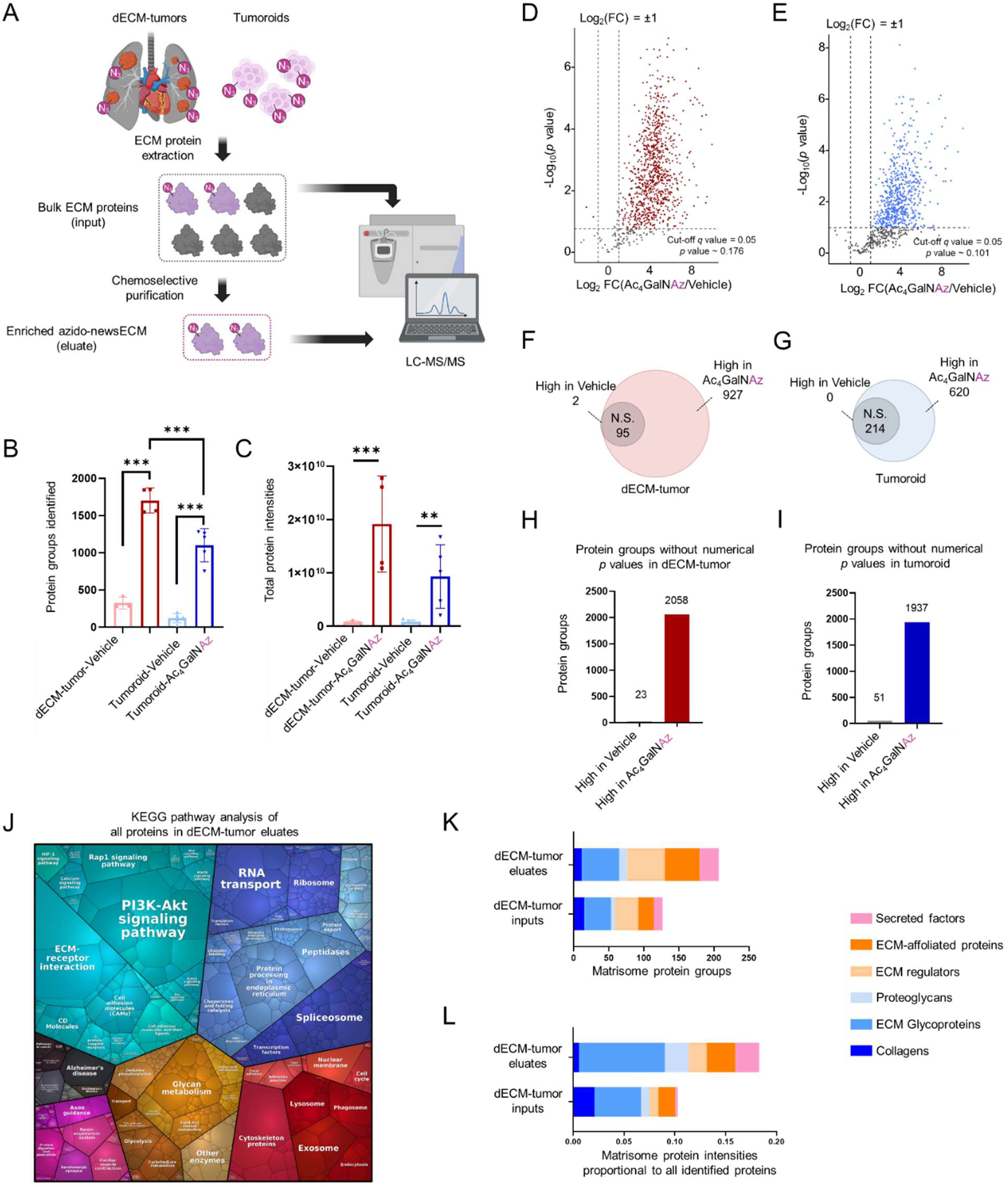
Targeted proteomics characterization and Gene Ontology (GO) analysis of newsECM profiles in bioengineered tumor models. (A) Schematic of the workflow for proteomics characterization of newsECM in dECM-tumors and tumoroids. (B) A bar graph plotting the protein groups identified in eluate samples from dECM-tumors receiving Ac_4_GalNAz (*n*=5) or DMSO (vehicle control, *n*=3) or tumoroids receiving Ac_4_GalNAz (*n*=5) or DMSO (vehicle control, *n*=5). C) A bar graph plotting the total protein intensities (before normalization) identified in eluate samples from dECM-tumors receiving Ac_4_GalNAz (*n*=4) or DMSO (vehicle control, *n*=3) or tumoroids receiving Ac_4_GalNAz (*n*=5) or DMSO (vehicle control, *n*=5). (D-I) Differential comparisons of protein intensities between Ac_4_GalNAz and DMSO (vehicle control) eluates from dECM-tumors (D,F,H) or tumoroids (E,G,I). Protein groups with comparisons that cannot be assigned with numerical *p* values (due to having fewer than 2 nonzero values in either treatment group) were pre-filtered out and plotted as bar graphs based on their average intensities in each treatment group (H,I). (D) Volcano plot for proteins identified in eluate samples from dECM-tumors receiving Ac_4_GalNAz (*n*=4) or DMSO (vehicle control, *n*=3). The proteins fall into three categories: high in Ac_4_GalNAz (red, Log_2_FC>1, *q*<0.05), high in Vehicle (black, Log_2_FC<-1, *q*<0.05) and non-significant (N.S., gray, *q*≥0.05 or -1≤Log_2_FC≤1). (E) Volcano plot for proteins identified in eluate samples from tumoroids receiving Ac_4_GalNAz (*n*=5) or DMSO (vehicle control, *n*=5). The proteins fall into three categories: high in Ac_4_GalNAz (blue, Log_2_FC>1, *q*<0.05), high in Vehicle (black, Log_2_FC<-1, *q*<0.05) and non-significant (N.S., gray, *q*≥0.05 or -1≤Log_2_FC≤1). For both panel D and E, the decimal intensities were transformed into Log_2_ and analyzed with two-tailed *t*-test with Welch’s correction, followed by multiple hypothesis testing correction with permutation-based FDR (FDR < 0.05) to compute *q* values. FCs were computed as fold changes of geometric means. The *p* value cut-off lines were plotted based on *q*=0.05 (*p* ∼ 0.176 in panel D; *p* ∼ 0.101 in panel E). (F,G) Venn diagrams showing the distribution among the three categories in panel D or panel E. J) A KEGG pathway Proteomap generated with all eluate proteins from the dECM-tumor receiving Ac_4_GalNAz. The area of each pathway represents its enrichment level and the color-coding represents different categories. (K,L) Matrisome analysis of inputs and eluates from dECM-tumors. The numbers (K) or total intensities (L) of proteins identified for each matrisome category were plotted for both inputs and eluates of the dECM-tumors. *** *p*<0.001. \*\**p*<0.01. Data are presented as means ± SD. Schematics were created with Biorender.com and published with permission.

### Gene Ontology analysis of newsECM profiles from dECM-tumors

Upon validating newsECM enrichment at the proteomics level, we went on to investigate the newsECM protein deposition patterns starting with the dECM-tumors. We first enriched the top KEGG pathways from the dECM-tumor eluates based on the quantitative composition of each individual protein ^[38]^. This revealed that the most active KEGG pathways linked to newsECM production in the dECM-tumors are associated with PI3K-Akt signaling, ECM-receptor interaction, cell adhesion, RNA transport, and exosome (Figure 4J). This supports the biological relevance of the newsECM profiling, as the PI3K-Akt cascade and the pathways associated with cell-substrate adhesion are frequently overactivated in native tumor tissues ^[45]^. The abundance plot of individual proteins (Figure S4) also shows higher abundance of ECM structural proteins such as Laminins (e.g. LAMA3, LAMC2, LAMB3), Tenascin-C (TNC), Fibronectin 1 (FN1) and Heparan Sulfate Proteoglycan 2 (HSPG2), as well as ECM/cell surface signaling proteins such as Integrins (e.g. ITGB1, ITGA6, ITGB4), suggesting that the tumor cells are actively depositing new ECM components onto the pre-existing dECM scaffold to reinforce interactions with their extracellular environment. The gene ontology (GO) functional annotation clustering ^[39,40]^of the top 100 abundant proteins in newsECM from the dECM-tumors also enriched terms with high relevance, including “ECM-receptor interaction”, “proteoglycans in cancer” and “extracellular matrix” (Table 1). To further categorize the revealed matrisomal protein synthesis, we matched the proteins identified in newsECM (eluates) and bulk ECM (inputs) from the dECM-tumors to a human matrisome dataset ^[46]^. Our results suggest that compared to bulk proteomics, newsECM profiling identifies more matrisome protein groups in general, especially in ECM regulators (21 more), ECM-affiliated proteins (28 more) and secreted factors (14 more) (Figure 4K). We also plotted the matrisome protein intensities in proportion to all identified proteins in eluates or inputs and observed that while newsECM detects less scaffolding proteins like collagens (only 26% of those found in inputs), it identifies higher intensities of other matrisome proteins such as ECM regulators (2.0-fold of those found in inputs), proteoglycans (2.9-fold of those found in inputs), and secreted factors (9.2-fold of those found in inputs) (Figure 4L). These analyses suggest that our approach, by selectively profiling newsECM, avoids the high background signals generated by pre-existing structural proteins within the biomaterial scaffold and offers an augmented capability to detect low-abundance extracellular species involved in ECM remodeling.

**Table 1.** Functional annotation clustering of proteins with top 100 abundance from dECM-tumor newsECM.

**Table 2.** Individual protein intensities before or/and after normalization in all three searches.

### Differential analysis reveals distinct newsECM patterns regulated by pre-existing dECM

To compare the differences in ECM synthesis patterns of the tumor cells when cultured with or without pre-existing ECM, we compared the newsECM profiles between dECM-tumors and ECM-free tumoroids. The correlation dot plot shows high correlation between replicates within each tissue model and low correlation between the replicates from two different models (Figure 5A), suggesting high consistency of the newsECM profiling as well as distinct newsECM profiles influenced by the pre-existing dECM. This is further supported by scatter plots of individual protein intensities showing high correlation (R^2^ ∼ 0.94, Figure S5A,B) between replicates from the same tissue model and low correlation (R^2^ ∼ 0.49, Figure S5C) between replicates from different models. The volcano plot (Figure 5B) and Venn diagram (Figure 5C) show how many extracellular proteins have differential synthesis levels between the two models: 928 and 500 proteins with upregulated synthesis in dECM-tumors and ECM-free tumoroids, respectively. We enriched the GO “biological processes” terms for the top 100 abundant proteins from the newsECM in each model (Figure 5D,E) and identified generally similar top enriched GO terms for both tumor models, except for “cell-matrix adhesion” which was specifically highlighted in dECM-tumors. This suggests that the tumor cells can sense and adapt to their surrounding environment by tailoring their ECM secretome.

**Figure 5.**
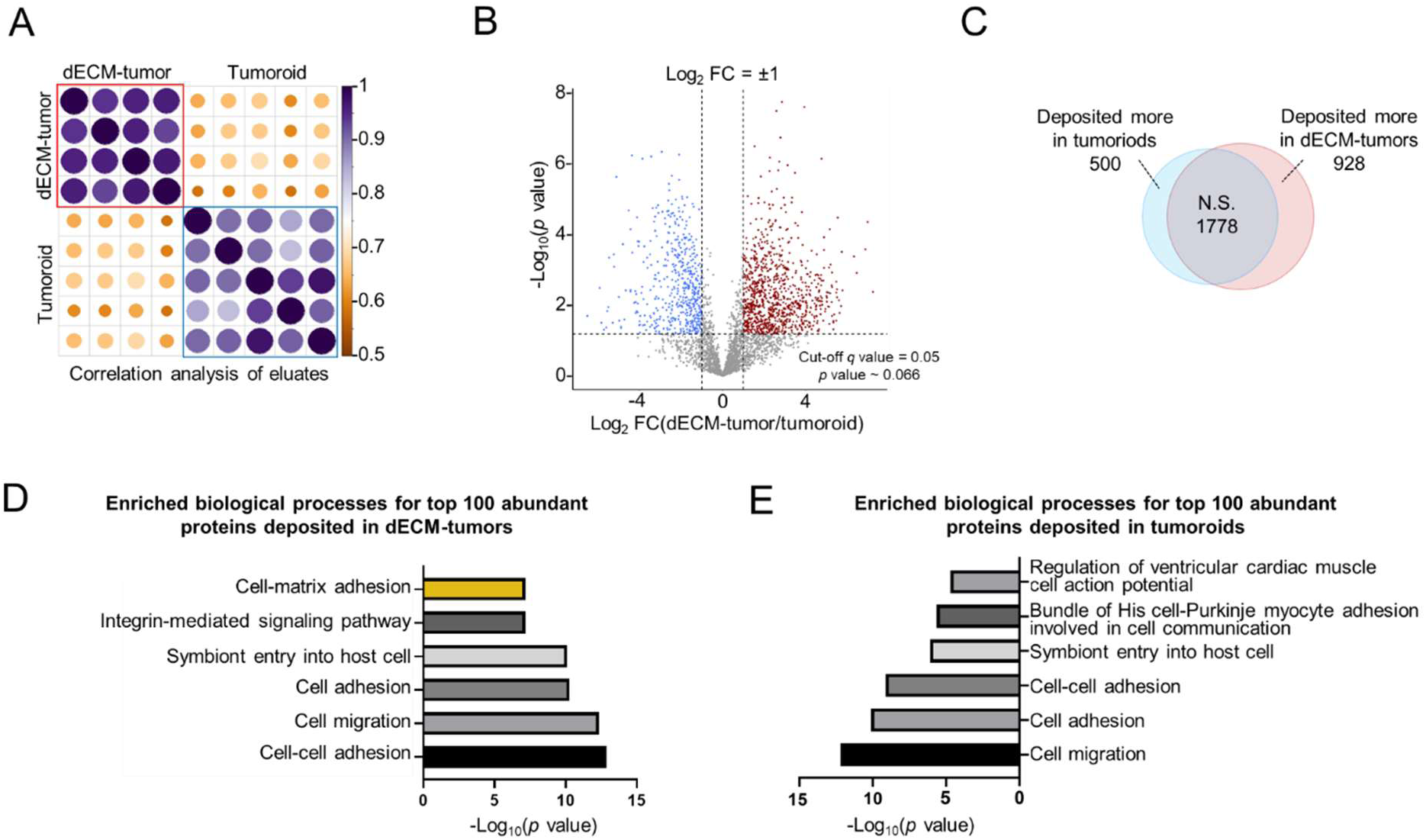
Differential analysis shows distinct newsECM patterns regulated by pre-existing dECM. (A) Correlation dot plot for proteomic profiles of eluates samples from dECM-tumors (*n*=4) and tumoroids (*n*=5) receiving Ac_4_GalNAz. Pearson correlation analysis was performed between each pair of samples. The colors indicate correlation levels, and the dot sizes indicate the significance of the correlation. Purple, high; Orange, low. Large, significant; Small, non-significant. (B) Volcano plot of proteins identified in eluate samples from dECM-tumors (*n*=4) or tumoroids (*n*=5) receiving Ac_4_GalNAz. The decimal intensities were transformed into Log_2_, imputed based on quantile regression imputation of left centered missing data (QRILC) method and analyzed with two-tailed *t*-test with Welch’s correction, followed by multiple hypothesis testing correction with permutation-based FDR (FDR < 0.05) to compute *q* values. The proteins fall into three categories: high in dECM tumors (red, Log_2_FC>1, *q*<0.05), high in tumoroids (blue, Log_2_FC<-1, *q*<0.05) and non-significant (N.S., gray, *q*≥0.05 or -1≤Log_2_FC≤1). FCs were computed as fold changes of geometric means. The *p* value cut-off line was plotted based on *q*=0.05 (*p* ∼ 0.066). (C) Venn diagram showing the distribution among three categories in panel B. (D,E) GO analysis of top enriched biological process terms of the 100 proteins with highest intensities in eluates samples from dECM-tumor (D) or tumoroid (E) receiving Ac_4_GalNAz.

### Elevated ECM digestive activities associated with pre-existing dECM

Tumor cells have long been recognized to actively degrade and remodel the ECM from the surrounding benign tissue environment to enhance tumor expansion and invasion ^[9]^. Laminins are crucial basement membrane components of the lung that play essential roles to support luminal epithelium and endothelium and to limit apical-to-basolateral cell migration ^[47]^. Intriguingly, immunofluorescence analysis of Laminin distribution in the dECM-tumor tissues revealed a reduction in LAMA1 abundance specifically within regions populated by the tumor nodes (Figure 1D). Inspired by this finding, we investigated the newsECM signatures for association with ECM digestive activities and identified Membrane-Type Matrix Metalloproteinase I (MT-MMP1, also known as MMP-14) with significantly elevated synthesis in dECM-tumors compared to ECM-free tumoroids (Figure 6A). This validated the biological relevance of our tissue models and newsECM profiling, as MMP-14 has reported activities to digest basement membrane components such as Collagens and Laminins ^[48–50]^, and its overexpression is frequently associated with cancer migration and metastasis ^[51,52]^. In parallel to eluates, we also searched the bulk proteomics results of input samples from both dECM-tumors and tumoroids, and were not able to detect MMP-14, further supporting the improved efficacy of newsECM profiling at identifying subtle changes in low-abundance ECM regulators compared to conventional proteomics. To validate this finding from the proteomics analysis, we performed immunofluorescence staining of MMP-14 in both tumor tissues and observed significantly elevated signals colocalizing with the cancer cells in the dECM-tumors in comparison to tumoroids (Figure 6B,C). This was further supported by MMP-14 western blot analysis on ECM extracts from both tumor models (Figure 6D,E, Figure S6). Together, our results demonstrated that tumor cells presented higher proteolytic and ECM-digestive activities when cultured with pre-existing dECM scaffolds.

**Figure 6.**
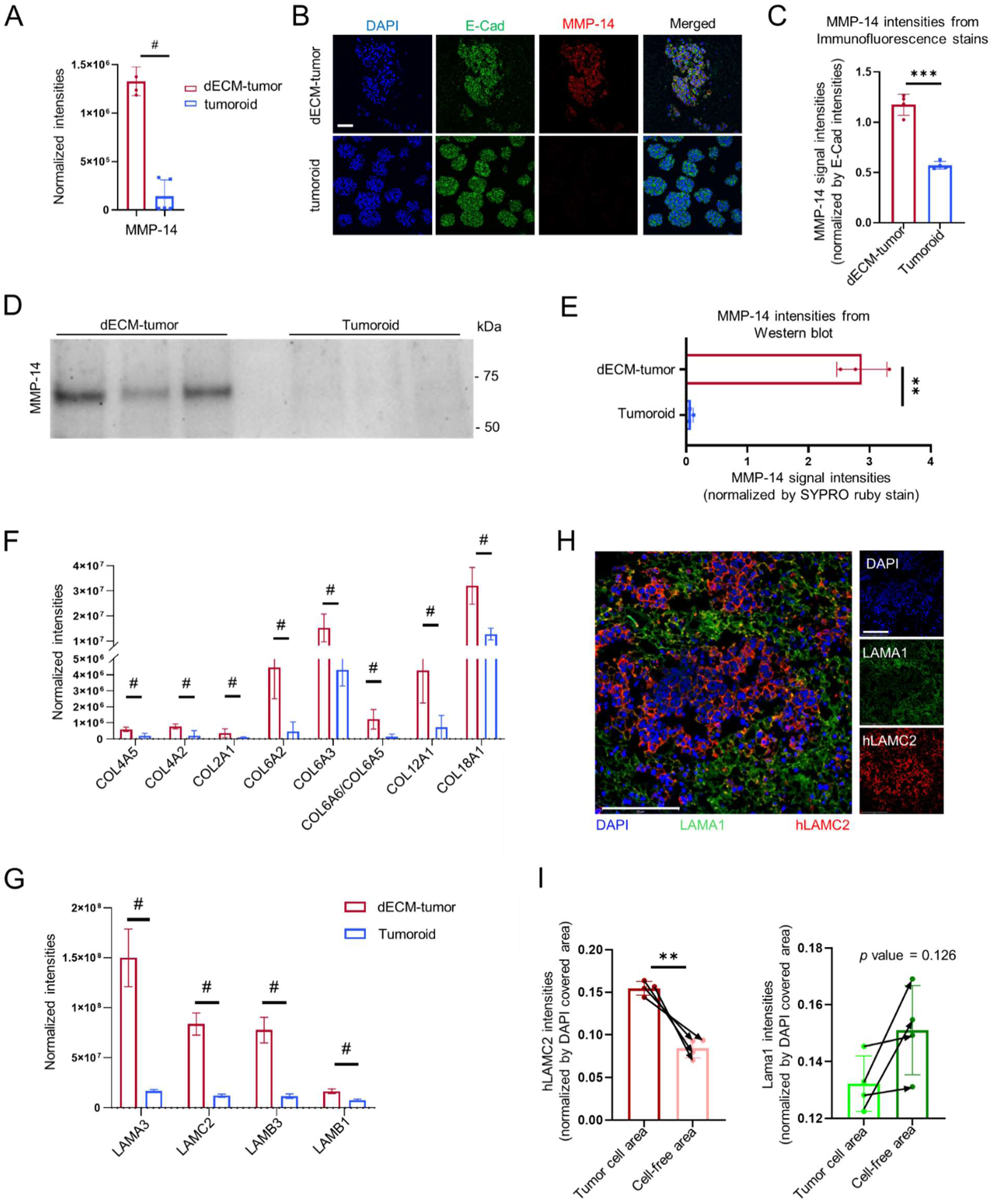
Tumor cells presented elevated remodeling activities when cultured with pre-existing ECM. (A) Bar graph showing the normalized, imputed protein intensities of MMP-14 in eluate samples from dECM-tumors (left, red, *n*=4) and tumoroids (right, blue, *n*=5) receiving Ac_4_GalNAz. The significance was determined by FDR-corrected *q* value (*q*=0.010 for MMP-14). (B) Immunofluorescence staining of MMP-14 (red) and E-Cad (green) in dECM-tumors (*n*=4) and tumoroids (*n*=4). Scale bar: 50 µm. (C) Quantification of MMP-14 signal intensities in areas populated with cells (based on DAPI signal) in panel B, normalized by E-Cad intensities. (D) Western blot detection of MMP-14 in the ECM fractions from dECM-tumors (left, *n*=3) or tumoroids (right, *n*=3). (E) Quantification of MMP-14 signal intensities in panel D, normalized by SYPRO Ruby dot blot of total proteins in Figure S6. (F,G) Bar graphs showing the normalized, imputed protein intensities of Collagen (F) or Laminin (G) subunits in eluate samples from dECM-tumors (red, *n*=4) or tumoroids (blue, *n*=5) receiving Ac_4_GalNAz. The significance was determined by FDR-corrected *q* values. (H) Immunofluorescence staining of LAMA1 (green) and hLAMC2 (red) in dECM-tumors (*n*=4). Scale bar: 275 µm. (I) Bar graphs quantifying LAMA1 and hLAMC2 signal intensities in panel H. Each image was separated into two parts based on DAPI signal: area populated with cells (with DAPI) and cell-free area (without DAPI). The hLAMC2 (left) and LAMA1 (right) signals on each area were plotted in the bar graphs. Arrows show data point connections for paired signal intensities from the same areas in the same images. ^#^ *q*<0.05; *** *p*<0.001; ** *p*<0.01. Data are presented as means ± SD. Zero values (when the protein was not observed by MS proteomics) were replaced with imputed values as described in Methods (Proteomic Data Analysis).

### Augmented tumor ECM deposition linked to pre-existing dECM

In parallel to aggressively digesting the surrounding benign tissue environments, tumor cells are well known to also actively lay down their own ECM species to build more favorable extracellular environments for their expansion and invasion ^[9]^. Comparing the newsECM profiles between the two tumor models, we observed augmented deposition of structural ECM components including several Collagen and Laminin subunits in dECM-tumors compared to tumoroids (Figure 6F,G). Notably, we identified significantly higher deposition levels of Laminin Alpha-3, Beta-3 and Gamma-2 subunits in dECM-tumors (Figure 6G), which are the subunits of Laminin-5 (Laminin-332) with reported roles in tumor progression and migration ^[53]^. To validate this finding, we performed immunofluorescence staining against tumor-associated Laminins (human Laminin Subunit Gamma 2, hLAMC2) in dECM-tumor sections and observed that hLAMC2 strongly colocalized with the tumor cell presence (Figure 6H,I left). Interestingly, we observed a reduction in LAMA1 (which is an important component of the basement membrane in healthy lungs^[54,55]^) at tumor cell loci (Figure 6H,I right), suggesting that the tumor cells within the dECM-tumors exert a synergistic influence to remodel their extracellular environment through combined ECM-digestive and synthetic activities. Taken together, our newsECM profiling establishes a comprehensive map documenting ECM deposition in bioengineered tumor models and how it is being modulated by pre-existing ECM environment or biomaterial support.

## Discussion

The ECM is a key constituent in nearly all native tissues and is accordingly being used extensively as scaffold material during tissue bioengineering for both disease modeling and regenerative medicine applications ^[11]^. Despite the wide recognition of the bidirectional nature of cell-ECM interactions, our understanding remains limited regarding how the biomaterial ECM regulates cells’ ability to remodel their extracellular environment. In this study, we developed a novel approach to chemoselectively profile the new ECM deposition by resident cells into their surrounding biomaterial ECM. We showcased the application of our analytical pipeline to two synthetic tumor tissue models with distinct extracellular environments. Comparing the newsECM proteomic profiles between the dECM-tumor and ECM-free tumoroid models, we uncovered a profound influence of pre-existing ECM on the ECM secretome of resident tumor cells. This observation was highlighted by elevated ECM remodeling mediated by augmented digestion of pre-existing ECM coupled with upregulated synthesis of tumor-associated ECM. We anticipate the technological platform described here to facilitate future inquiries regarding how cells respond to their surrounding biomaterial environments in a wide variety of tissue engineering scenarios as well as ECM remodeling events taking place in vivo.

Tumor cells are well-known for their ability to build an extracellular environment that favors their own growth and enhances their plasticity for metastasis ^[9]^. By comparing the newsECM profiles between two tumor models with and without foreign ECM support, we found that the presence of biomaterial scaffold derived from decellularized lung ECM directed the tumor cells to adopt an ECM deposition pattern that better recapitulates what is observed in native tumor tissues. For example, we detected with our untargeted newsECM proteomics analysis the augmented synthesis of MMP-14 in dECM-tumors, which has been reported as the driving force behind ECM destruction during tumor cell invasion ^[48,49,51,52]^. Protein-protein interaction network functional enrichment analysis with the STRING database showed that MMP-14 physically interacts with other critical ECM degrading enzymes such as MMP-2 and MMP-9 (Figure S7). In fact, MMP-14 is required for the processing and activation of MMP-2 ^[56]^ that is involved in degrading basement membrane proteins such as Collagen-IV ^[57]^. Additionally, a recent clinical study correlated upregulated MMP-14 with reduced Prospero Homeobox Protein 1 (PROX-1) ^[58]^, which is identified as a transcriptional repressor of MMP-14 ^[59]^. Interestingly, reduced PROX-1 synthesis was also captured in dECM-tumors compared to tumoroids in our study (Figure S8A). In addition to MMP-14, other ECM regulators, such as Procollagen C-Endopeptidase Enhancer 2 (PCOLCE2), a predictive marker for epithelial-to-mesenchymal transition and metastasis in lung cancer ^[60]^, were also observed with elevated synthesis levels in the dECM-tumors (Figure S8B). In line with lower ECM digestive activities in the ECM-free tumoroids, we observed a higher synthesis level of Tissue Inhibitor of Metalloproteinase 1 (TIMP-1), an MMP-2 inhibitor ^[61]^, in the tumoroids compared to dECM-tumors (Figure S8C).

Other than ECM proteins associated with matrix degradation, we also observed augmented deposition of tumor-associated basement membrane components in dECM-tumors compared to tumoroids. For example, in dECM-tumors we observed augmented synthesis of all three subunits of Laminin-332 (Figure 6G,H) that is closely associated with aggressive tumor invasion ^[53,62,63]^, with the Gamma 2 subunit being reported as a therapeutic target against tumor aggravation ^[64]^. In addition to Laminins, in dECM-tumors we also observed elevated deposition of Alpha-2 Chain of Type IV Collagen (COL4A2, Figure 6G), another major component of the tumor microenvironment with potential involvement in the pathogenesis and progression of multiple cancer types, including lung cancer ^[65]^. Collectively, our results suggest that the presence of a dECM scaffold tunes the tumor ECM deposition towards patterns with improved resemblance to those found in native tumors, and that investigating the ECM secretomes of the cells can be an effective way to assess pathophysiological relevance of bioengineered tissue models.

Glycosylation is a common PTM with high occurrence in proteins secreted to the extracellular space (predicted to be over 90% in humans ^[23–25]^). Therefore, bioorthogonal labeling via glycosylation has the potential of tagging a significant portion of newly synthesized ECM proteins. Our chemoselective newsECM profiling approach enriches extracellular proteins newly deposited by resident cells over a defined period, free from the pre-existing ones of the biomaterial scaffold, enabling their selective characterization with improved detection sensitivity. In addition, our approach also avoids the proteomics background generated by high-abundance structural ECM proteins (such as Collagens) and allows more focused detection of ECM species with signaling and regulatory roles which are often of low abundances. For example, we were able to identify MMP-14 with high consistency in the dECM-tumor eluates with newsECM profiling (Figure 6A) but were not able to detect any MMP-14 in the inputs with conventional bulk proteomics. In addition, the relatively low turnover rates of structural ECM proteins lead to their accumulation over time on bioengineered tissues ^[66]^, thus masking the detection with regular mass spectrometry of signaling proteins that undergo more dynamic turnover. The described newsECM profiling offers a strategy to overcome this challenge and enables study of the dynamics of ECM regulators that may be under-investigated by traditional proteomics. Consistent with our study, a recent report pioneered the labeling and visualization of nascent matrix proteins produced by chondrocytes within polymer hydrogels by directly targeting new peptide synthesis using an azido-bearing methionine analog (L-azidohomoalanine) ^[67]^. This study together with ours highlights the versatility of click chemistry-based strategies for probing cellular remodeling of their surrounding matrix environment.

Beyond deposition of new ECM structural components, effective degradation of pre-existing ECM is frequently observed during tissue morphogenesis, repair and regeneration to enable regulated cell migration, invasion, and expansion. Similarly, tumor cells have a well-documented ability to degrade benign tissue ECM to favor their expansion and metastasis. Accordingly, investigating the cell-ECM dynamics should encompass the understanding of coordinated constructive and digestive remodeling by resident cells on their surrounding extracellular environments. The newsECM profiling presented here, together with several other relevant technologies such as SILAC ^[16]^ and bioorthogonal non-canonical amino acid tagging (BONCAT) ^[67,68]^, all have a direct focus on characterizing new protein synthesis. Interestingly, with the revealed upregulation of MMP-14 in dECM-tumors and the correlated finding of selective degradation of pre-existing basement membrane proteins, we demonstrate that the characterization of newsECM can also offer important insights into proteome-wide degradation events, proving a more comprehensive view of ECM remodeling and turnover.

We envision that the utility of the newsECM profiling pipeline described herein can be extended beyond synthetic tumor tissues, as it can serve as a platform technology applicable to nearly all cell-incorporated, tissue-engineered systems to reveal fundamental mechanisms of cell-ECM interaction. As demonstrated in the ECM-free tumoroid model, we also anticipate our approach to facilitate studies in material-free organoid or spheroid cultures to reveal temporally regulated ECM production in response to induced changes in genetic, environmental, or therapeutic conditions. Building upon our prior study demonstrating the feasibility of effective labeling of newsECM in a wide variety of tissues and organs in vivo ^[26,29]^, we anticipate the chemoselective proteomic pipeline described here to be applicable to in vivo animal models and open enormous opportunities for novel mechanistic insight regarding coordinated ECM remodeling mediating organogenesis and pathogenesis. Lastly, leveraging new developments in ex vivo tissue perfusion and culture (such as precision-cut tissue slice) ^[69,70]^ and our recent demonstration of newly synthesized glycoprotein labeling during ex vivo lung perfusion ^[27]^, our approach also opens the door to direct inquiry into ECM dynamics in human tissues bearing specific pathophysiological conditions.

## Supporting Information

**Supplementary figure 1.** Western blot detection of azido→biotin signal in cellular fractions of dECM-tumors.

**Supplementary figure 2.** Western blot detection of azido→biotin signal in cellular fractions of tumoroids.

**Supplementary figure 3.** Bar graph of human or rat protein intensities in eluate versus input from the dECM-tumors.

**Supplementary figure 4.** An individual-protein-intensity Proteomap generated with all eluate proteins from the dECM-tumor receiving Ac_4_GalNAz.

**Supplementary figure 5.** Scatter plots of normalized protein intensities in eluate samples from dECM-tumors or tumoroids.

**Supplementary figure 6.** SYPRO Ruby dot blot of total proteins from each sample analyzed in Figure 6D.

**Supplementary figure 7.** Protein-protein physical interaction network functional enrichment analysis with the STRING database.

**Supplementary figure 8.** Bar graphs showing the normalized, imputed protein intensities of (A) PROX-1, (B) PCOLCE2 and (C) TIMP-1 in eluate samples from dECM-tumors (left, red, *n*=4) and tumoroids (right, blue, *n*=5).

## Acknowledgements

We thank Nicholas Keuler from the Statistical Consulting Group at the University of Wisconsin – Madison for providing advice on statistical analysis. We thank Biospecimen Core at the University of Pittsburgh for histological processing, the Vivarium at Carnegie Mellon University for animal husbandry, and Misti West and Garrett Struble for laboratory management.

## Data Availability

Data will be made available upon reasonable request.

## References

[1] Y. Xing, B. Varghese, Z. Ling, A. S. Kar, E. Reinoso Jacome, X. Ren, Regen Eng Transl Med 2022, 8, 55.

[2] C. Frantz, K. M. Stewart, V. M. Weaver, J Cell Sci 2010, 123, 4195.

[3] C. Bonnans, J. Chou, Z. Werb, Nat Rev Mol Cell Biol 2014, 15, 786.

[4] T. Rozario, D. W. DeSimone, Dev Biol 2010, 341, 126.

[5] A. Naba, Nature Reviews Molecular Cell Biology 2024 25:11 2024, 25, 865.

[6] D. P. Tabassum, K. Polyak, Nature Reviews Cancer *2015 15:8* 2015, 15, 473.

[7] S. Zhang, X. Xiao, Y. Yi, X. Wang, L. Zhu, Y. Shen, D. Lin, C. Wu, Signal Transduction and Targeted Therapy *2024 9:1* 2024, 9, 1.

[8] J. J. F. Sleeboom, G. S. van Tienderen, K. Schenke-Layland, L. J. W. van der Laan, A. A. Khalil, M. M. A. Verstegen, Sci Transl Med 2024, 16, 3840.

[9] J. Winkler, A. Abisoye-Ogunniyan, K. J. Metcalf, Z. Werb, Nat Commun 2020, 11, 5120.

[10] K. S. Hwang, E. U. Seo, N. Choi, J. Kim, H. N. Kim, Bioact Mater 2023, 21, 576.

[11] G. Jensen, C. Morrill, Y. Huang, Acta Pharm Sin B 2018, 8, 756.

[12] E. E. Wheeler, J. K. Leach, Tissue Eng Part B Rev 2024, DOI 10.1089/TEN.TEB.2024.0212.

[13] J. J. Song, H. C. Ott, Trends Mol Med 2011, 17, 424.

[14] Z. Zhao, X. Chen, A. M. Dowbaj, A. Sljukic, K. Bratlie, L. Lin, E. L. S. Fong, G. M. Balachander, Z. Chen, A. Soragni, M. Huch, Y. A. Zeng, Q. Wang, H. Yu, Nature Reviews Methods Primers *2022 2:1* 2022, 2, 1.

[15] P. Wijesekara, P. Yadav, L. A. Perkins, D. B. Stolz, J. M. Franks, S. C. Watkins, E. Reinoso Jacome, S. L. Brody, A. Horani, J. Xu, A. Barati Farimani, X. Ren, iScience 2022, 25, DOI 10.1016/J.ISCI.2022.104730.

[16] S. E. Ong, B. Blagoev, I. Kratchmarova, D. B. Kristensen, H. Steen, A. Pandey, M. Mann, Molecular & Cellular Proteomics 2002, 1, 376.

[17] O. Rosmark, E. Åhrman, C. Müller, L. Elowsson Rendin, L. Eriksson, A. Malmström, O. Hallgren, A. K. Larsson-Callerfelt, G. Westergren-Thorsson, J. Malmström, Scientific Reports *2018 8:1* 2018, 8, 1.

[18] S. Garbis, G. Lubec, M. Fountoulakis, J Chromatogr A 2005, 1077, 1.

[19] Z. Wang, M. Gerstein, M. Snyder, Nature Reviews Genetics *2008 10:1* 2009, 10, 57.

[20] D. Greenbaum, C. Colangelo, K. Williams, M. Gerstein, Genome Biol 2003, 4, DOI 10.1186/GB-2003-4-9-117.

[21] S. P. Gygi, Y. Rochon, B. R. Franza, R. Aebersold, Mol Cell Biol 1999, 19, 1720.

[22] J. Eichler, Curr Biol 2019, 29, R229.

[23] M. Lynch, J. Barallobre-Barreiro, M. Jahangiri, M. Mayr, J Intern Med 2016, 280, 325.

[24] H. J. An, J. W. Froehlich, C. B. Lebrilla, Curr Opin Chem Biol 2009, 13, 421.

[25] K. J. Colley, A. Varki, R. S. Haltiwanger, T. Kinoshita, Essentials of Glycobiology 2022, DOI 10.1101/GLYCOBIOLOGY.4E.4.

[26] X. Ren, D. Evangelista-Leite, T. Wu, K. T. Rajab, P. T. Moser, K. Kitano, K. P. Economopoulos, D. E. Gorman, J. P. Bloom, J. J. Tan, S. E. Gilpin, H. Zhou, D. J. Mathisen, H. C. Ott, Biomaterials 2018, 182, 127.

[27] Z. Ling, K. Noda, B. L. Frey, M. Hu, S. W. Fok, L. M. Smith, P. G. Sanchez, X. Ren, Am J Physiol Lung Cell Mol Physiol 2023, 325, L30L44.

[28] Y. Xing, S. S. Yerneni, W. Wang, R. E. Taylor, P. G. Campbell, X. Ren, Biomaterials 2022, 281, DOI 10.1016/J.BIOMATERIALS.2021.121357.

[29] Z. Ling, Y. Xing, E. R. Jacome, S. W. Fok, X. Ren, Bio Protoc 2021, 11, DOI 10.21769/BIOPROTOC.3922.

[30] L. F. Tapias, S. E. Gilpin, X. Ren, L. Wei, B. C. Fuchs, K. K. Tanabe, M. Lanuti, H. C. Ott, Ann Thorac Surg 2015, 100, 414.

[31] V. V. Rostovtsev, L. G. Green, V. V. Fokin, K. B. Sharpless, Angewandte Chemie International Edition 2002, 41, 2596.

[32] J. Schindelin, I. Arganda-Carreras, E. Frise, V. Kaynig, M. Longair, T. Pietzsch, S. Preibisch, C. Rueden, S. Saalfeld, B. Schmid, J. Y. Tinevez, D. J. White, V. Hartenstein, K. Eliceiri, P. Tomancak, A. Cardona, Nature Methods *2012 9:7* 2012, 9, 676.

[33] C. S. Hughes, S. Moggridge, T. Müller, P. H. Sorensen, G. B. Morin, J. Krijgsveld, Nature Protocols *2018 14:1* 2018, 14, 68.

[34] S. K. Solntsev, M. R. Shortreed, B. L. Frey, L. M. Smith, J Proteome Res 2018, 17, 1844.

[35] S. Tyanova, T. Temu, P. Sinitcyn, A. Carlson, M. Y. Hein, T. Geiger, M. Mann, J. Cox, Nature Methods *2016 13:9* 2016, 13, 731.

[36] S. H. Yu, D. Ferretti, J. P. Schessner, J. D. Rudolph, G. H. H. Borner, J. Cox, Curr Protoc Bioinformatics 2020, 71, DOI 10.1002/CPBI.105.

[37] C. Lazar, T. Burger, CRAN: Contributed Packages 2022, DOI 10.32614/CRAN.PACKAGE.IMPUTELCMD.

[38] W. Liebermeister, E. Noor, A. Flamholz, D. Davidi, J. Bernhardt, R. Milo, Proc Natl Acad Sci U S A 2014, 111, 8488.

[39] D. W. Huang, B. T. Sherman, R. A. Lempicki, Nat Protoc 2009, 4, 44.

[40] B. T. Sherman, M. Hao, J. Qiu, X. Jiao, M. W. Baseler, H. C. Lane, T. Imamichi, W. Chang, Nucleic Acids Res 2022, 50, W216.

[41] D. Szklarczyk, R. Kirsch, M. Koutrouli, K. Nastou, F. Mehryary, R. Hachilif, A. L. Gable, T. Fang, N. T. Doncheva, S. Pyysalo, P. Bork, L. J. Jensen, C. Von Mering, Nucleic Acids Res 2023, 51, D638.

[42] S. Gunti, A. T. K. Hoke, K. P. Vu, N. R. London, Cancers (Basel) 2021, 13, 874.

[43] X. Ren, P. T. Moser, S. E. Gilpin, T. Okamoto, T. Wu, L. F. Tapias, F. E. Mercier, L. Xiong, R. Ghawi, D. T. Scadden, D. J. Mathisen, H. C. Ott, Nature Biotechnology *2015 33:10* 2015, 33, 1097.

[44] J. D. Hirsch, L. Eslamizar, B. J. Filanoski, N. Malekzadeh, R. P. Haugland, J. M. Beechem, R. P. Haugland, Anal Biochem 2002, 308, 343.

[45] M. J. Sanaei, S. Razi, A. Pourbagheri-Sigaroodi, D. Bashash, Transl Oncol 2022, 18, 101364.

[46] A. Naba, K. R. Clauser, S. Hoersch, H. Liu, S. A. Carr, R. O. Hynes, Molecular and Cellular Proteomics 2012, 11, DOI 10.1074/mcp.M111.014647.

[47] L. Schuger, Exp Lung Res 1997, 23, 119.

[48] K. L. Sodek, M. J. Ringuette, T. J. Brown, British Journal of Cancer *2007 97:3* 2007, 97, 358.

[49] J. Ueda, M. Kajita, N. Suenaga, K. Fujii, M. Seiki, Oncogene *2003 22:54* 2003, 22, 8716.

[50] E. L. Bair, L. C. Man, K. McDaniel, K. Sekiguchi, A. E. Cress, R. B. Nagle, G. T. Bowden, Neoplasia 2005, 7, 380.

[51] E. M. Wenzel, N. M. Pedersen, L. A. Elfmark, L. Wang, I. Kjos, E. Stang, L. Malerød, A. Brech, H. Stenmark, C. Raiborg, Nature Communications *2024 15:1* 2024, 15, 1.

[52] A. M. Knapinska, G. B. Fields, Pharmaceuticals 2019, 12, 77.

[53] Y. Moriya, T. Niki, T. Yamada, Y. Matsuno, H. Kondo, S. Hirohashi, 2001, DOI 10.1002/1097-0142(20010315)91:6.

[54] G. Burgstaller, B. Oehrle, M. Gerckens, E. S. White, H. B. Schiller, O. Eickelberg, Eur Respir J 2017, 50, DOI 10.1183/13993003.01805-2016.

[55] N. M. Nguyen, R. M. Senior, Dev Biol 2006, 294, 271.

[56] K. Y. Han, J. Dugas-Ford, M. Seiki, J. H. Chang, D. T. Azar, Invest Ophthalmol Vis Sci 2015, 56, 5323.

[57] S. Monaco, V. Sparano, M. Gioia, D. Sbardella, D. Di Pierro, S. Marini, M. Coletta, Protein Sci 2006, 15, 2805.

[58] L. Claesson-Welsh, J Clin Invest 2020, 130, 1093.

[59] S. Gramolelli, J. Cheng, I. Martinez-Corral, M. Vähä-Koskela, E. Elbasani, E. Kaivanto, V. Rantanen, K. Tuohinto, S. Hautaniemi, M. Bower, C. Haglund, K. Alitalo, T. Mäkinen, T. V. Petrova, K. Lehti, P. M. Ojala, Sci Rep 2018, 8, DOI 10.1038/S41598-018-27739-W.

[60] S. Bin Lim, S. J. Tan, W. T. Lim, C. T. Lim, Nature Communications *2017 8:1* 2017, 8, 1.

[61] V. Arpino, M. Brock, S. E. Gill, Matrix Biology 2015, 44–46, 247.

[62] P. Rousselle, J. Y. Scoazec, Semin Cancer Biol 2020, 62, 149.

[63] B. G. Kim, H. J. An, S. Kang, Y. P. Choi, M. Q. Gao, H. Park, N. H. Cho, Am J Pathol 2011, 178, 373.

[64] M. Garg, G. Braunstein, H. P. Koeffler, Expert Opin Ther Targets 2014, 18, 979.

[65] H. JingSong, G. Hong, J. Yang, Z. Duo, F. Li, C. WeiCai, L. XueYing, M. YouSheng, O. Y. YiWen, P. Yue, C. Zou, Oncotarget 2016, 8, 2585.

[66] Q. Li, Z. Chang, G. Oliveira, M. Xiong, L. M. Smith, B. L. Frey, N. V. Welham, Biomaterials 2015, 81, 104.

[67] C. Loebel, A. M. Saleh, K. R. Jacobson, R. Daniels, R. L. Mauck, S. Calve, J. A. Burdick, Nature Protocols *2022 17:3* 2022, 17, 618.

[68] D. C. Dieterich, A. J. Link, J. Graumann, D. A. Tirrell, E. M. Schuman, Proc Natl Acad Sci U S A 2006, 103, 9482.

[69] T. Cruz, A. L. Mora, M. Rojas, Methods Mol Biol 2021, 2299, 139.

[70] H. N. Alsafadi, F. E. Uhl, R. H. Pineda, K. E. Bailey, M. Rojas, D. E. Wagner, M. Königshoff, Am J Respir Cell Mol Biol 2020, 62, 681.

