## Supporting Information for "Chemoselective Characterization of New Extracellular Matrix Deposition in Bioengineered Tumor Tissues"

### Supplementary Data

**Supplementary figure 1.** Western blot detection of azido→biotin signal in cellular fractions of dECM-tumors.

**Supplementary figure 2.** Western blot detection of azido→biotin signal in cellular fractions of tumoroids.

**Supplementary figure 3.** Bar graph of human or rat protein intensities in eluate versus input from the dECM-tumors.

**Supplementary figure 4.** An individual-protein-intensity Proteomap generated with all eluate proteins from the dECM-tumor receiving $Ac_4GalNAz$.

**Supplementary figure 5.** Scatter plots of normalized protein intensities in eluate samples from dECM-tumors or tumoroids.

**Supplementary figure 6.** SYPRO Ruby dot blot of total proteins from each sample analyzed in Figure 6D.

**Supplementary figure 7.** Protein-protein physical interaction network functional enrichment analysis with the STRING database.

**Supplementary figure 8.** Bar graphs showing the normalized, imputed protein intensities of (A) PROX-1, (B) PCOLCE2 and (C) TIMP-1 in eluate samples from dECM-tumors (left, red, $n$=4) and tumoroids (right, blue, $n$=5).

**Table 1.** Functional annotation clustering of proteins with top 100 abundance from dECM-tumor newsECM.

**Table 2.** Individual protein intensities before or/and after normalization in all three searches. (Separate excel file)

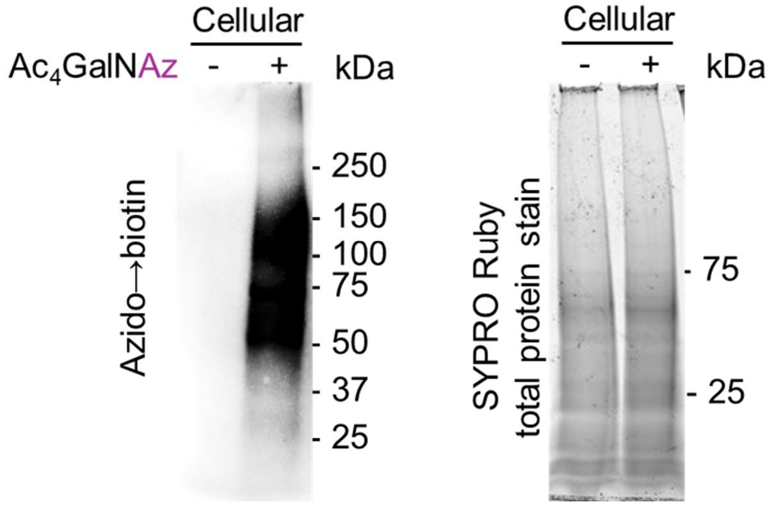


**Supplementary figure 1. Western blot detection of azido→biotin signal in cellular fractions of dECM-tumors.** Western blot detection of azido→biotin signal in the cellular fractions of dECM-tumors using streptavidin-HRP (left) and SYPRO Ruby staining of total proteins (right).

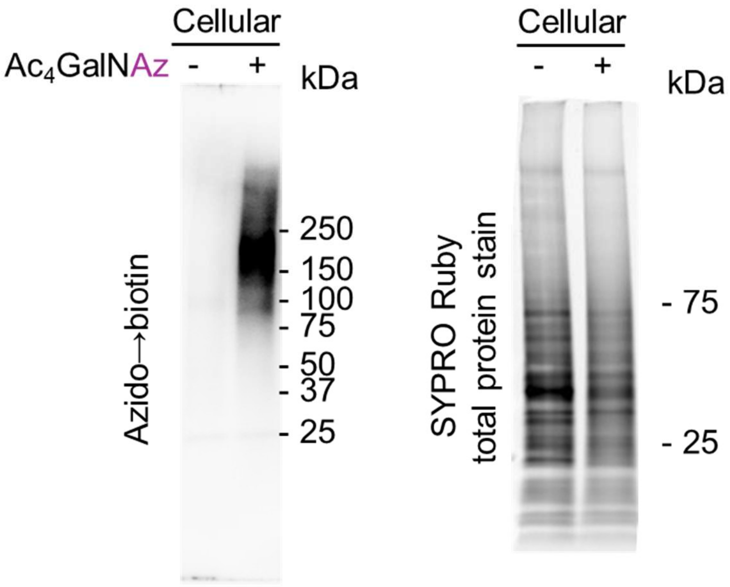


**Supplementary figure 2. Western blot detection of azido→biotin signal in cellular fractions of tumoroids.** Western blot detection of azido→biotin signal in the cellular fractions of tumoroids using streptavidin-HRP (left) and SYPRO Ruby staining of total proteins (right).

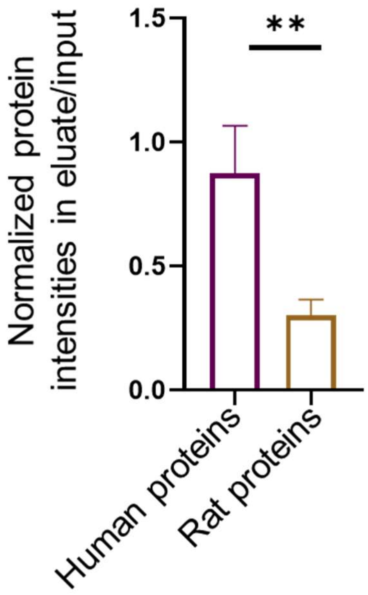


**Supplementary figure 3. Bar graph of human or rat protein intensities in eluate versus input from the dECM-tumors.** $n$=4. ** $p$<0.01. Data are presented as means ± SD.

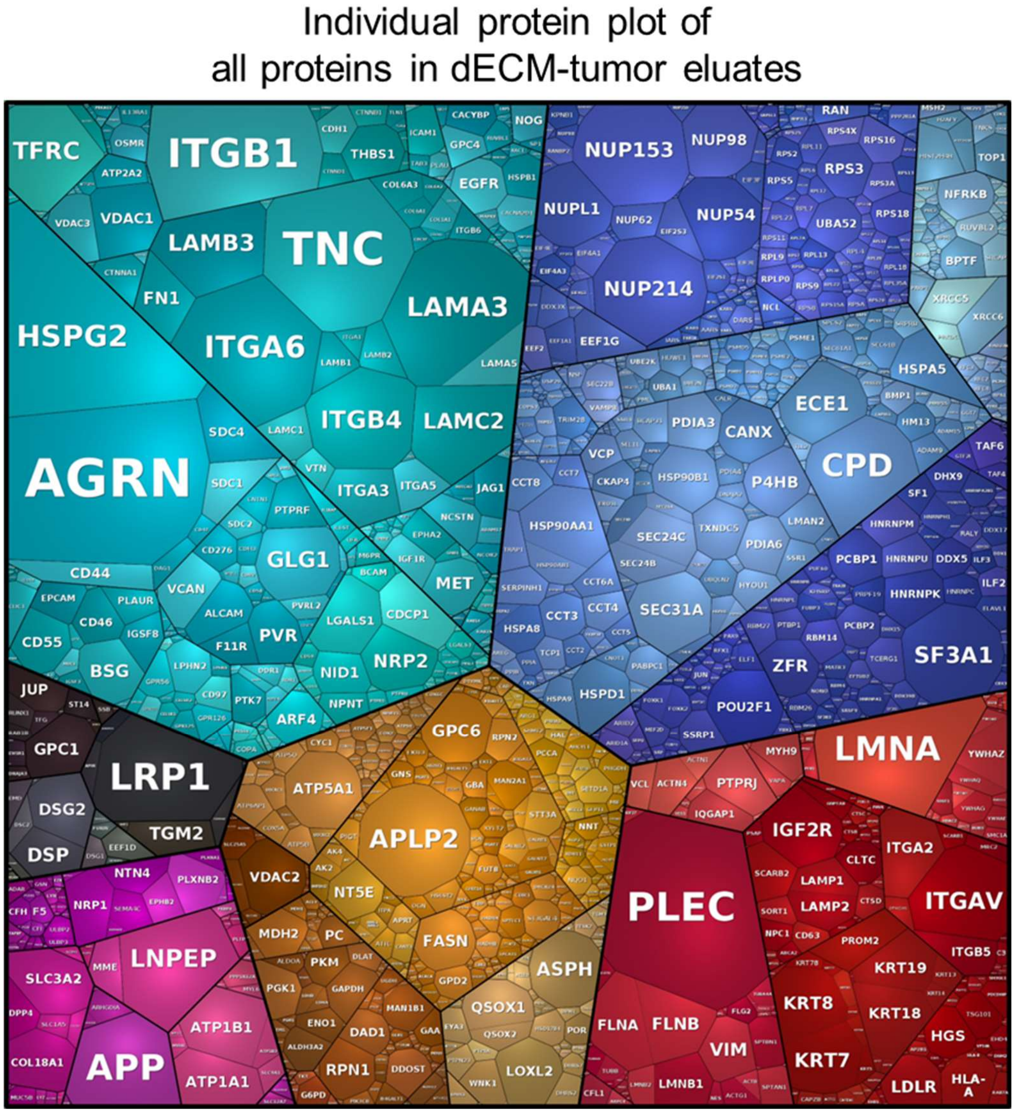


**Supplementary figure 4. An individual-protein-intensity Proteomap generated with all eluate proteins from the dECM-tumor receiving $Ac_4GalNAz$.** The area of each protein represents its intensity level and color-coded for different proteins.

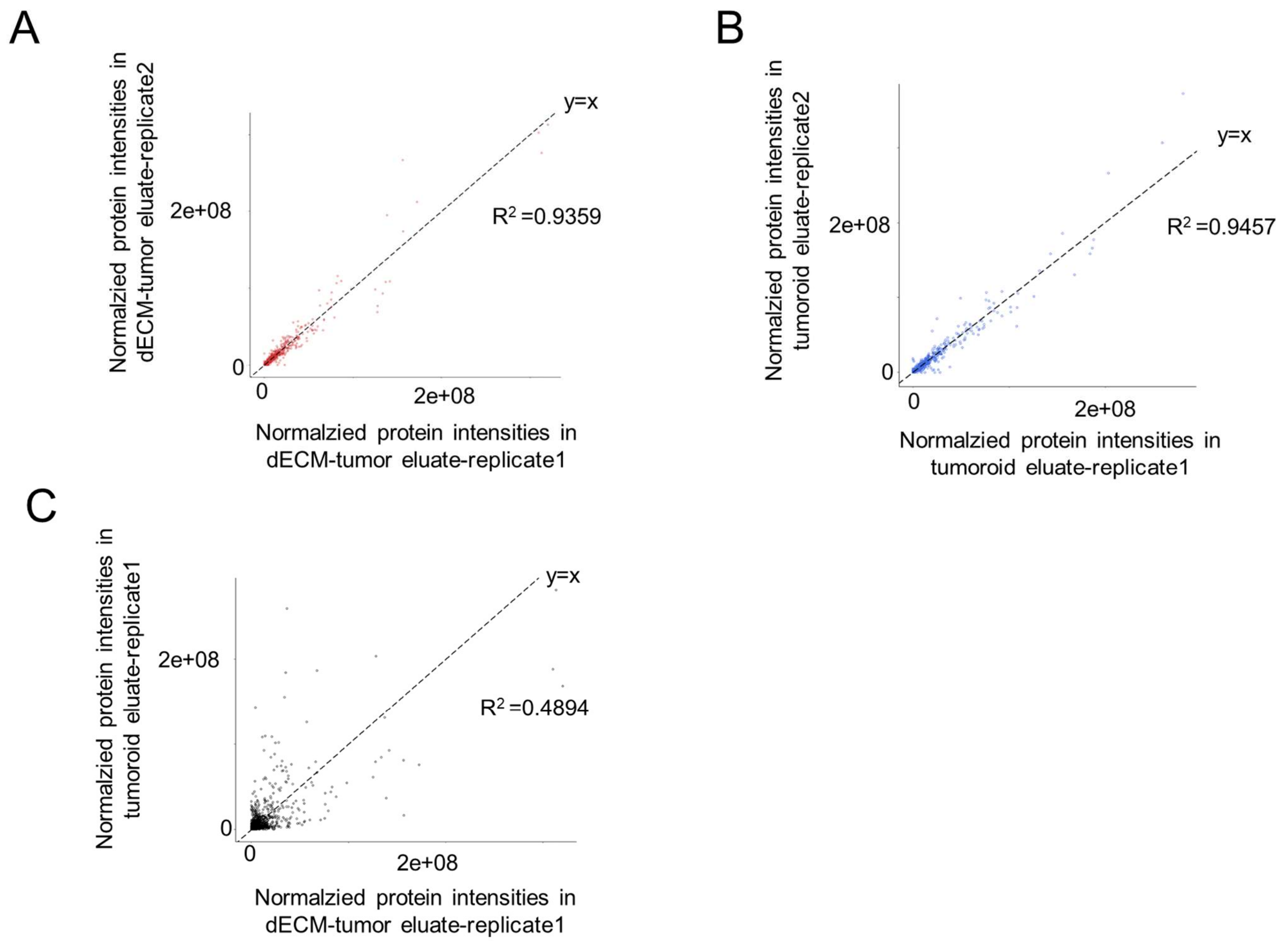


**Supplementary figure 5. Scatter plots of normalized protein intensities in eluate samples from dECM-tumors or tumoroids.** Scatter plots of protein intensities between (A) two eluate samples from dECM-tumor, (B) two eluate samples from tumoroids, and (C) one eluate sample from dECM-tumor and one eluate sample from tumoroid.

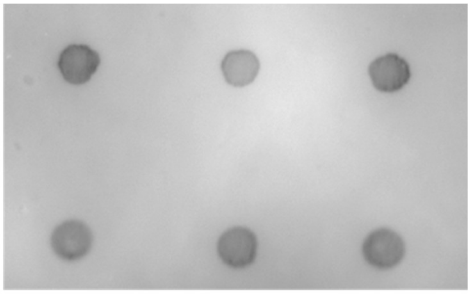


**Supplementary figure 6. SYPRO Ruby dot blot of total proteins from each sample analyzed in Figure 6D.**

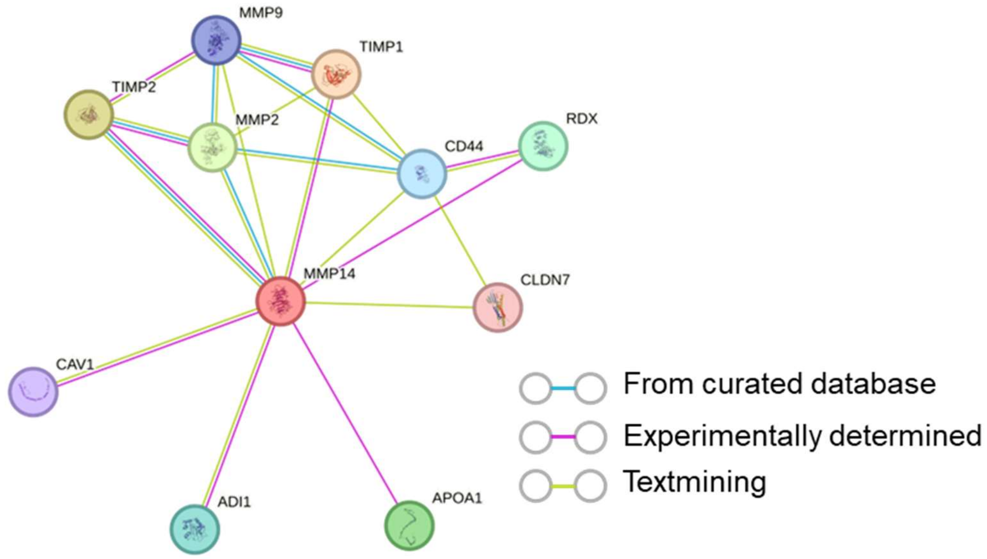


**Supplementary figure 7. Protein-protein physical interaction network functional enrichment analysis with the STRING database.** The colors of the lines linking two proteins represent the information sources of known or predicted physical interactions.

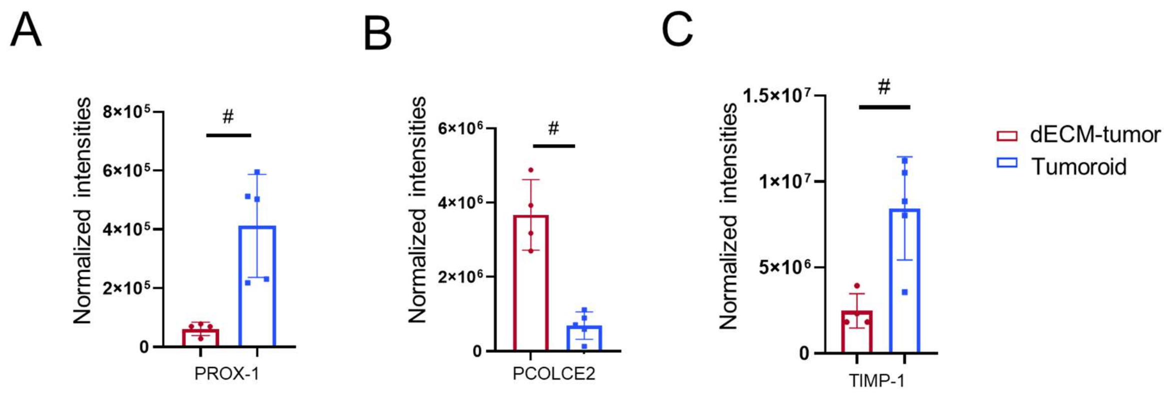


**Supplementary figure 8. Bar graphs showing the normalized, imputed protein intensities of (A) PROX-1, (B) PCOLCE2 and (C) TIMP-1 in eluate samples from dECM-tumors (left, red, *n*=4) and tumoroids (right, blue, *n*=5).** # $q$<0.05. Data are presented as means ± SD. Zero values (when the protein was not observed by MS proteomics) were replaced with imputed values as described in Methods (Proteomic Data Analysis).

**Table 1. Functional annotation clustering of proteins with top 100 abundance from dECM-tumor newsECM.**

| GO category | GO term | Count | *p* value | Adjusted *p* value[1] |
|---|---|---|---|---|
| Annotation Cluster 1 (Enrichment score: 12.41) | | | | |
| KEGG_PATHWAY | ECM-receptor interaction | 18 | 1.90E-19 | 2.60E-17 |
| GOTERM_BP_DIRECT | Cell migration | 17 | 4.40E-13 | 2.50E-10 |
| KEGG_PATHWAY | Proteoglycans in cancer | 12 | 6.70E-07 | 1.50E-05 |
| Annotation Cluster 2 (Enrichment score: 10.48) | | | | |
| GOTERM_CC_DIRECT | Collagen-containing extracellular matrix | 21 | 6.40E-15 | 5.40E-13 |
| GOTERM_CC_DIRECT | Basement membrane | 10 | 1.00E-09 | 3.10E-08 |
| UP_KW_CELLULAR_COMPONENT | Extracellular matrix | 14 | 5.70E-09 | 6.00E-08 |
| Annotation Cluster 3 (Enrichment score: 9.61) | | | | |
| UP_SEQ_FEATURE | CARBOHYD:N-linked (GlcNAc...) asparagine | 57 | 9.10E-14 | 1.20E-10 |
| UP_KW_DOMAIN | Signal | 59 | 2.50E-10 | 5.40E-09 |
| UP_KW_PTM | Disulfide bond | 52 | 6.70E-07 | 4.50E-06 |
| Annotation Cluster 4 (Enrichment Score: 8.99) | | | | |
| UP_KW_PTM | Proteoglycan | 14 | 1.70E-13 | 2.30E-12 |
| UP_KW_PTM | Heparan sulfate | 7 | 5.40E-09 | 4.90E-08 |
| GOTERM_CC_DIRECT | lysosomal lumen | 9 | 2.50E-08 | 5.90E-07 |
| GOTERM_CC_DIRECT | Golgi lumen | 9 | 5.00E-08 | 1.00E-06 |

[1] *p* values were adjusted by Benjamini correction with FDR<0.05.

**Table 2. Individual protein intensities before or/and after normalization in all three searches.** "*Search 1* original": original 17 eluates (1 outliner removed) files that underwent GPTMD and were searched using the Human XML database with MBR. "*Search 1* normalization 1": for comparison between $Ac_4GalNAz$ and Vehicle groups, the protein intensities were normalized separately in each treatment group. "*Search 1* normalization 2": proteins from 9 $Ac_4GalNAz$ dECM-tumor and tumoroid eluate samples were normalized all together for their proportional intensities in each group. "*Search 2* original": original 15 dECM-tumor files (inputs and eluates, 1 eluate removed) that underwent GPTMD analysis and were searched using the Human and Rat XML databases without MBR. "*Search 3* original": original 18 dECM-tumor and tumoroid input files that underwent GPTMD analysis and were searched using the Human and Rat XML databases with MBR. "*Search 3* normalization": all input samples were normalized together. The normalization formulas can be found under "Proteomic Data Analysis" in Method section. The table contents are in a separate excel file.
